# Fluoroquinolone resistance-conferring *gyrA* variants alter the fitness cost and potentiate the resistance of the zoliflodacin resistance mutation *gyrB*^D429N^ in *Neisseria gonorrhoeae*

**DOI:** 10.64898/2026.05.04.722797

**Authors:** Aditi Mukherjee, Sofia OP Blomqvist, David Helekal, Daniel HF Rubin, Bailey Bowcutt, Samantha G Palace, Yonatan H Grad

**Affiliations:** Department of Immunology and Infectious Diseases, Harvard T. H. Chan School of Public Health, Boston, Massachusetts, USA

**Author notes:** **Corresponding author’s contact information:** Yonatan Grad, Department of Immunology and Infectious Diseases, Harvard T.H. Chan School of Public Health, Boston, MA 02115, USA.

## Abstract

*Neisseria gonorrhoeae* is a major public health concern due to its high global prevalence and rapid evolution of antibiotic resistance. A first-in-class topoisomerase inhibitor, zoliflodacin (a spiropyrimidinetrione) recently received FDA approval for treatment of gonorrhea, but its potential for cross-resistance with another topoisomerase inhibitor, the fluoroquinolone antibiotic ciprofloxacin, remains poorly understood. Here, we investigated how genetic diversity in the fluoroquinolone target *gyrA* influences the resistance and fitness effects of the zoliflodacin resistance mutation *gyrB*^D429N^. We constructed an isogenic panel of *N. gonorrhoeae* to determine how the resistance and fitness effects of the *gyrB*^D429N^ mutation are modulated by the most common ciprofloxacin resistance-associated variants in *gyrA*. In the presence of *gyrB*^D429N^, the zoliflodacin minimum inhibitory concentration (MIC) was 2-4-fold higher in strains that also contained ciprofloxacin resistance-associated *gyrA* alleles, and the *gyrB*^D429N^ mutation reciprocally increased ciprofloxacin MICs of these strains 3-6-fold. Fitness cost of the *gyrB*^D429N^ mutation varied from modest to severe across *gyrA* backgrounds, with the largest cost in ciprofloxacin resistant *gyrA*^91F/95G^ and *gyrA*^91F/95N^ backgrounds and comparatively minimal cost in the ciprofloxacin resistant *gyrA*^91F/95A^ background. These results demonstrate the capacity for epistatic interactions among resistance-associated *gyrA* and *gyrB* mutations, underscoring the need for genomic surveillance to monitor high-risk combinations of resistance determinants as new therapies are deployed.

## Introduction

*Neisseria gonorrhoeae*, the etiologic agent of gonorrhea, remains a major global public health challenge and a priority pathogen for the World Health Organization due to its high prevalence and rising rates of antimicrobial resistance (AMR)^1^. Historically, sulfonamides, penicillin, tetracycline, and fluoroquinolones served as first-line therapies for gonorrhea, but, as resistance emerged to each antibiotic, the rate of clinical failures rendered them inappropriate for empiric therapy^2^. Currently, ceftriaxone is the sole remaining antibiotic recommended for first-line, empiric treatment of gonorrhea^3^, though two new, first-in-class antibiotics, zoliflodacin and gepotidacin, recently received FDA approval for urogenital gonorrhea^4^. Given the limited treatment options and the expectation that resistance will emerge, understanding how resistance mutations interact across drug classes can guide strategies for optimizing use of therapeutics and for monitoring the emergence of resistance.

Fluoroquinolones provide an example of the evolutionary plasticity that enables rapid development of resistance in *N. gonorrhoeae*. Resistance to ciprofloxacin emerged through substitutions in the quinolone resistance-determining regions of DNA gyrase and topoisomerase IV, most commonly involving amino acid changes at positions 91 and 95 of *gyrA* and 86-91 in *parC*^5^. Strains carrying these mutations spread rapidly and have shaped global gonococcal population structure, with the prevalence of ciprofloxacin resistance in much of Asia at or near 100%^6^ and ∼30-40% in the US^7^.

Recent advances in gonorrhea treatment have introduced two novel first-in-class topoisomerase inhibitors with mechanisms distinct from those of fluoroquinolones. Zoliflodacin targets the B subunit of DNA gyrase, and gepotidacin inhibits both DNA gyrase and topoisomerase IV. Both drugs demonstrated clinical efficacy against ciprofloxacin-resistant *N. gonorrhoeae*^8, 9^. Resistance to zoliflodacin can arise through substitutions in *gyrB*, including D429N, K450N, and K450T^13, 14^. For gepotidacin, resistance arose during the phase 2 trial via spontaneous *gyrA*^A92T^ mutation in two isolates that harbored *parC*^D86N^, a mutation also associated with ciprofloxacin resistance^8, 15–17^.

Zoliflodacin and gepotidacin were developed to be effective against strains with established fluoroquinolone-resistance mechanisms. However, their deployment into populations already enriched for topoisomerase mutations raises the possibility that resistance to these new drugs may arise in strains with pre-existing ciprofloxacin resistance alleles. Interactions between these resistance alleles may modulate both drug susceptibility and evolutionary outcomes. Building on the observation that ciprofloxacin resistance-associated substitutions in *parC* also contribute to gepotidacin resistance, recent work suggests that additional interactions among ciprofloxacin, gepotidacin, and zoliflodacin resistance variants may facilitate cross-resistance and influence strain fitness^13, 18^. However, critical questions remain about the interactions among resistance-associated loci for these drugs. In particular, ciprofloxacin use has driven diversity in *gyrA* in *N. gonorrhoeae*^19, 20^. Use of the new topoisomerase-targeting antimicrobials will select for resistance on the background of this *gyrA* diversity. Will zoliflodacin resistance via the *gyrB*^D429N^ mutation be more beneficial in certain strain backgrounds, as might be expected if *gyrB*^D429N^ collaterally increases ciprofloxacin resistance^13^ or if the degree of zoliflodacin resistance conferred by *gyrB*^D429N^ is influenced by *gyrA* alleles? How does epistasis between *gyrB*^D429N^ and *gyrA* modulate resistance-associated fitness costs to influence in which strains *gyrB*^D429N^ is more likely to arise and spread?

In this study, we evaluated the epistatic interactions among zoliflodacin and ciprofloxacin resistance mutations using a panel of isogenic strains carrying common *gyrA* alleles with and without the *gyrB*^D429N^ mutation. For each strain, we determined the antimicrobial susceptibility to four topoisomerase inhibitors: ciprofloxacin, zoliflodacin, gepotidacin, and nalidixic acid. We then quantified the *in vitro* fitness effect of zoliflodacin resistance in the context of each *gyrA* allele using pairwise fitness assays and amplicon sequencing.

## Results

### The extent of zoliflodacin and ciprofloxacin resistance conferred by *gyrB*^D429N^ is modulated by *gyrA* alleles

Acquisition of the zoliflodacin-resistance mutation *gyrB*^D429N^ can increase resistance to ciprofloxacin in some, but not all, backgrounds^13, 21^. To probe the genetic variation that might influence cross-resistance, we evaluated how susceptibility to topoisomerase-targeting drugs is modulated by the interaction of *gyrB*^D429N^ with common ciprofloxacin resistance-associated *gyrA* alleles. We constructed a panel of isogenic strains in the *N. gonorrhoeae* clinical isolate GCGS0481, a strain background in which *gyrB*^D429N^ can increase ciprofloxacin resistance^13^. The panel was composed of isogenic strains with mutations at *gyrA* positions 91 (*gyrA*^91S^, associated with ciprofloxacin susceptibility, or *gyrA*^91F^, associated with ciprofloxacin resistance) and 95 (the ciprofloxacin susceptible *gyrA*^95D^ allele or the variants *gyrA*^95G/A/N^ associated with increases in ciprofloxacin resistance) in all pairwise combinations. We then introduced the zoliflodacin resistance mutation *gyrB*^D429N^ to each strain and measured zoliflodacin, ciprofloxacin, gepotidacin, and nalidixic acid MICs (**Table 1**).

**Table 1:**
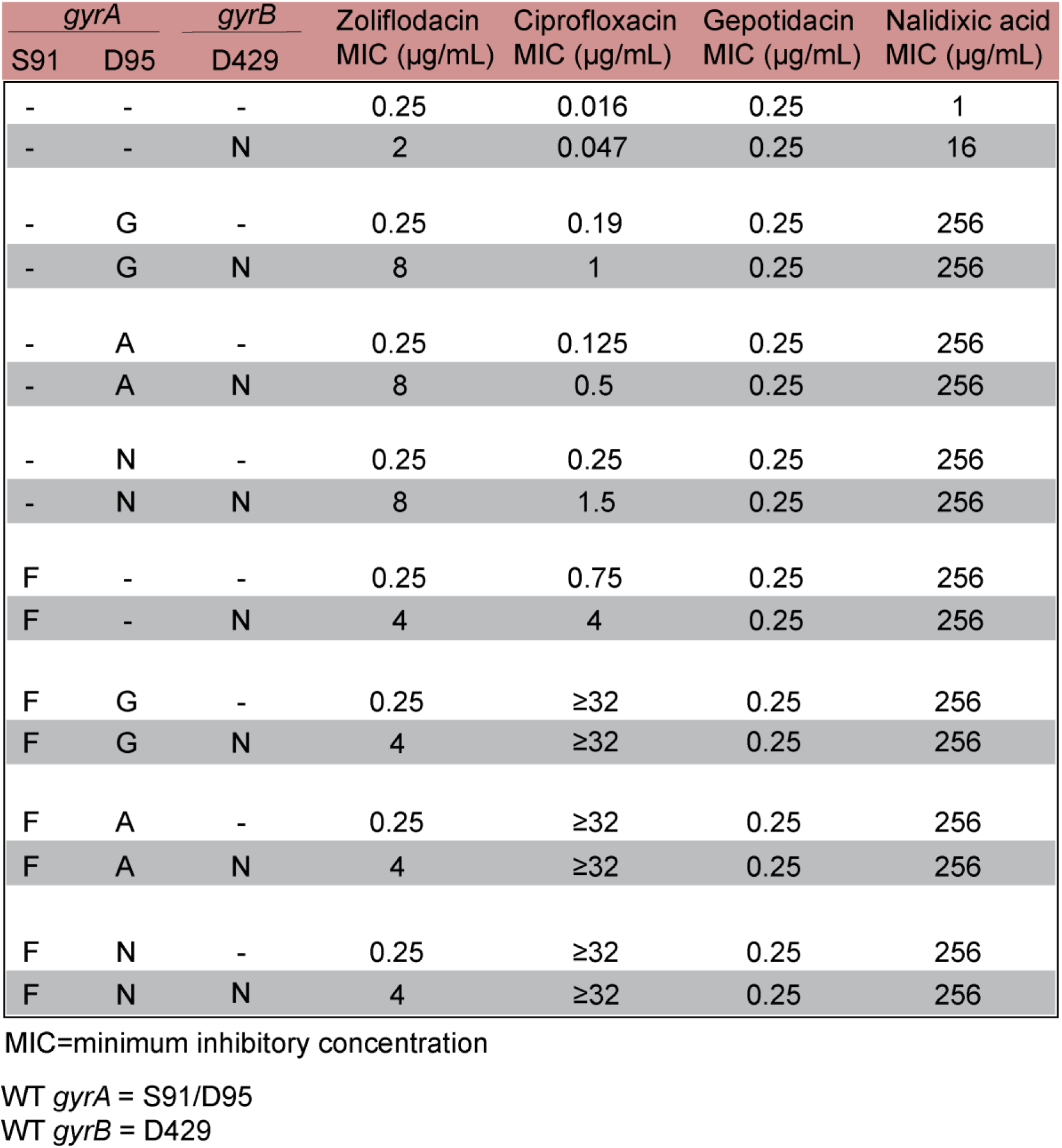
Zoliflodacin, ciprofloxacin, gepotidacin, and nalidixic acid MICs of isogenic *gyrA* and *gyrB* variants in the *N. gonorrhoeae* GCGS0481 background. Hyphens denote that the strain contains the reference allele at the given position.

As expected, MICs corresponding to clinical ciprofloxacin resistance (MIC ≥ 1 μg/mL) were exclusively found in mutants carrying the *gyrA*^91F^ mutation, and none of the *gyrA* alleles tested altered zoliflodacin susceptibility in the absence of the *gyrB*^D429N^ mutation (**Table 1**). The extent of zoliflodacin resistance conferred by *gyrB*^D429N^ varied by *gyrA* allele. Zoliflodacin MICs were 2-to-4-fold higher in backgrounds carrying ciprofloxacin resistance-associated mutations in *gyrA* relative to the baseline ciprofloxacin susceptible allele *gyrA*^91S/95D^ (**Table 1**). The *gyrB*^D429N^ mutation also led to an increase in ciprofloxacin MICs across multiple mutant *gyrA* backgrounds compared to the isogenic *gyrB*^429D^ controls: *gyrB*^D429N^ increased ciprofloxacin MIC by ∼3-fold in the ciprofloxacin susceptible (*gyrA*^91S/95D^) background and 4-to-6-fold in strains carrying one ciprofloxacin resistance-associated variant at *gyrA* position 91 or 95 (*gyrA*^91S/95G^, *gyrA*^91S/95A^, *gyrA*^91S/95N^, *gyrA*^91F/95D^). Strains with variants at both *gyrA* positions 91 and 95 have high-level ciprofloxacin resistance (MICs ≥32 µg/mL), which was maintained in the presence of *gyrB*^D429N^. In addition to its role in zoliflodacin resistance, *gyrB*^D429N^ contributes to nalidixic acid resistance in *N. gonorrhoeae*^22–25^. Consistent with this expectation, *gyrB*^D429N^ increased the nalidixic acid MIC of the susceptible *gyrA*^91S/95D^ strain 16-fold (from 1 to 16 μg/mL). Ciprofloxacin resistance-associated mutations also confer nalidixic acid resistance^26, 27^, demonstrated by the 256-fold increase observed in all mutant *gyrA* strains in the panel (from 1 to 256 µg/mL). However, unlike ciprofloxacin, no difference in susceptibility was observed for nalidixic acid resistance in various mutant *gyrA* backgrounds with or without *gyrB*^D429N^ (**Table 1**). In all strains, gepotidacin MICs remained unchanged.

### Relative fitness of zoliflodacin-resistant *gyrB*^D429N^ strains is associated with *gyrA* allele

We hypothesized that zoliflodacin resistance-associated fitness costs may be modulated by *gyrA* genotype. We therefore determined the relative fitness of *gyrB*^D429N^ mutants derived from each *gyrA* variant tested above. We first evaluated the *in vitro* growth pattern of each isogenic *gyrA* variant strain and its *gyrB*^D429N^ derivative in antibiotic-free gonococcal medium in monoculture. We observed no significant differences in growth among the *gyrA* strains without additional *gyrB* mutation (**Supplementary Figure 1**), but in the presence of *gyrB*^D429N^, strains carrying the *gyrA*^91F/95N^ and *gyrA*^91F/95G^ mutations had reduced growth (**Supplementary Figure 2**).

We performed competition assays to determine the relative fitness of each strain in the panel compared to the baseline, drug susceptible strain (*gyrA*^91S/95D^ *gyrB*^429D^). First, we employed pairwise competition assays, in which two strains that differ by an antibiotic resistance marker are co-cultured, their relative abundance is measured by differential plating, and relative fitness is inferred from the net change in strain frequencies over time. We measured the relative abundance of mutant strains over 8 hours of co-culture. Among ciprofloxacin resistant strains carrying *gyrA*^91F^, the fitness cost of the zoliflodacin-resistant *gyrB*^D429N^ derivatives was comparable in the background with no additional *gyrA*^95^ mutation (*gyrA*^91F/95D^) and in the *gyrA*^91F/95A^ background (**Figure 1A, C**), but more severe in the *gyrA*^91F/95G^ and *gyrA*^91F/95N^ backgrounds (**Figure 1B, D**). In contrast to *gyrA*^91F^ backgrounds, pairwise competition assays were largely unable to detect fitness defects associated with *gyrB*^D429N^ in the ciprofloxacin-susceptible *gyrA*^91S/95D^ background or in most of its corresponding *gyrA*^95^ variants (**Figure 1E, G, H**). A detectable cost emerged only in the *gyrA*^91S/95G^ background at 8 hours, the final timepoint of the assay (**Figure 1F**), suggesting that fitness differences in this background accumulated slowly and became apparent only as cultures approached stationary phase. These results were generally in agreement with the slower growth observed for the *gyrA*^91F/95G^ *gyrB*^D429N^ and *gyrA*^91F/95N^ *gyrB*^D429N^ strains in monoculture (**Supplementary Figure 2**), with the additional sensitivity of the competition method detecting fitness costs for the *gyrA*^91F/95D^ *gyrB*^D429N^ and *gyrA*^91F/95A^ *gyrB*^D429N^ strains. All pairwise competition results were consistent with the reciprocal competition in which the kanamycin marker was carried by the opposite strain (**Supplementary Figure 3A-H**).

**Figure 1:**
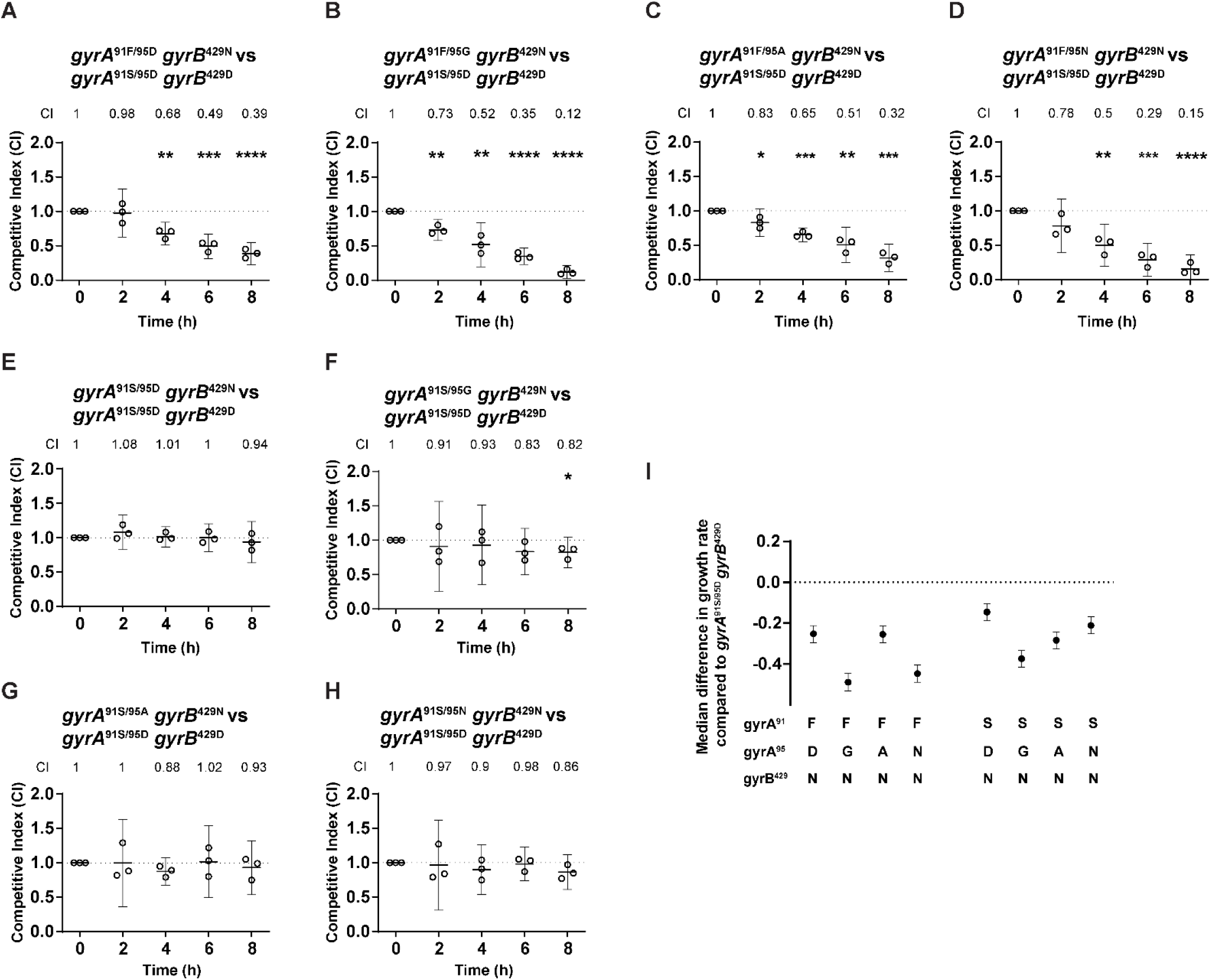
Relative fitness of the *gyrB*^D429N^ mutants in isogenic GCGS0481 strains with various *gyrA* alleles. Y-axes show the competitive index (CI) (panels A-H) or fitness difference (panel I) of each *gyrB*^D429N^ mutant relative to the ciprofloxacin-susceptible strain *gyrA*^91S/95D^ *gyrB*^429D^. In pairwise assays (panel A to H), the *gyrB*^D429N^ mutant carried a kanamycin marker and was cocultured with its unmarked parental strain; shown: mean with 95% confidence interval, with statistically significant differences in competitive indices compared to time 0 analyzed using an unpaired Student’s *t-test*, indicated *p ≤ 0.05, **p ≤ 0.01, ***p ≤ 0.001, and ****p ≤ 0.0001. n = 3, representative of three independent experiments performed in absence of any antibiotic pressure. (A) *gyrA*^91F/95D^ *gyrB*^429N^: p = 0.91, 0.25, 0.01, 0.005, respectively for 2, 4, 6, and 8 hours. (B) *gyrA*^91F/95G^ *gyrB*^429N^: p = 0.24, 0.99, 0.046, 0.03, respectively for 2, 4, 6, and 8 hours. (C) *gyrA*^91F/95A^ *gyrB*^429N^: p = 0.24, 0.29, 0.0003, 0.005, respectively for 2, 4, 6, and 8 hours. (D) *gyrA*^91F/95N^ *gyrB*^429N^: p = 0.69, 0.25, 0.01, 0.004, respectively for 2, 4, 6, and 8 hours. (E) *gyrA*^91S/95D^ *gyrB*^429N^: p = 0.62, 0.13, 0.14, 0.09, respectively for 2, 4, 6, and 8 hours. (F) *gyrA*^91S/95G^ *gyrB*^429N^: p = 0.1, 0.21, 0.04, 0.75, respectively for 2, 4, 6, and 8 hours. (G) *gyrA*^91S/95A^ *gyrB*^429N^: p = 0.48, 0.99, 0.2, 0.47, respectively for 2, 4, 6, and 8 hours. (H) *gyrA*^91S/95N^ *gyrB*^429N^: p = 0.23, 0.1, 0.05, 0.22, respectively for 2, 4, 6, and 8 hours. (I) Estimated difference in exponential-phase growth rate between *gyrA*^91S/95D^ *gyrB*^429D^ and each mutant strain, calculated from pooled competition experiments via amplicon sequencing of barcodes; Shown: median and 95% credible interval (CrI) estimate for difference in growth rate, estimated from 2 biological replicates of all barcoded strains grown together (4 barcodes per strain per replicate; timepoints at 0, 2, and 4 hours of coculture).

To enable direct comparisons of fitness across strains, we also measured competitive fitness using an amplicon sequencing based method. In contrast to the differential plating competition assays described above for paired strains, all strains were marked with both kanamycin and neutral nucleotide barcodes and co-cultured in a single large pool. The relative abundance of each was determined by amplicon sequencing of the barcode locus (**Methods**). This assay enabled relative fitness estimation of multiple strains from a single experiment, increasing sensitivity by reducing experimental noise. To avoid drop-out of especially unfit strains in this complex pool and to focus on relative fitness in exponential growth, we synchronized strains into log-phase growth prior to competition and restricted analysis to early timepoints.

As seen with pairwise competition, barcode amplicon sequencing identified a substantial cost of *gyrB*^D429N^ in ciprofloxacin resistant (*gyrA*^91F^) strains, with more extreme costs in *gyrA*^91F/95N^ and *gyrA*^91F/95G^ backgrounds and more moderate costs in *gyrA*^91F/95D^ and *gyrA*^91F/95A^ backgrounds (**Figure 1I**). The concordance between assays for these genotypes indicates that the fitness cost of *gyrB*^D429N^ in the *gyrA*^91F/95G^ and *gyrA*^91F/95N^ backgrounds originated during exponential growth and accumulated as cultures transitioned into stationary phase. Unlike differential plating competition assays, the amplicon sequencing method detected moderate but consistent fitness disadvantages of *gyrB*^D429N^ in all *gyrA*^91S^ backgrounds. This discrepancy likely reflects the higher sensitivity of the amplicon sequencing-based assay to small changes in exponential-phase growth rate.

### *gyrA* variants modulate the fitness cost of the *gyrB*^D429N^ mutation

The fitness effects presented above were measured relative to the ciprofloxacin and zoliflodacin susceptible strain GCGS0481 *gyrA*^91S/95D^ *gyrB*^429D^. We performed additional competition experiments to determine whether the observed fitness costs are attributable to the *gyrB*^D429N^ mutation itself, the *gyrA* alleles, or interactions among them. We first compared the fitness of each *gyrB*^D429N^ mutant directly to its isogenic *gyrA* parental strain. In pairwise competition assays, *gyrB*^D429N^ imposed a substantial fitness disadvantage in the *gyrA*^91F/95G^ and *gyrA*^91F/95N^ backgrounds (**Figure 2B, D**). In contrast, the fitness effects of *gyrB*^D429N^ were comparable and more moderate in the *gyrA*^91F/95D^ and *gyrA*^91F/95A^ backgrounds (**Figure 2A, C**). These same trends were observed via amplicon sequencing of barcoded strains in the pooled competition experiment (**Figure 2H**). *gyrB*^D429N^ did not confer a detectable fitness disadvantage in pairwise competitions in the ciprofloxacin-susceptible *gyrA*^91S^ background, regardless of the *gyrA*^95^ variant present (**Figure 2F–H**), although amplicon sequencing identified modest growth defects conferred by *gyrB*^D429N^ in these strains (**Figure 2H**). All pairwise competition results were consistent with the reciprocal competition in which the kanamycin marker was carried by the opposite strain (**Supplementary Figure 4A-H**).

**Figure 2:**
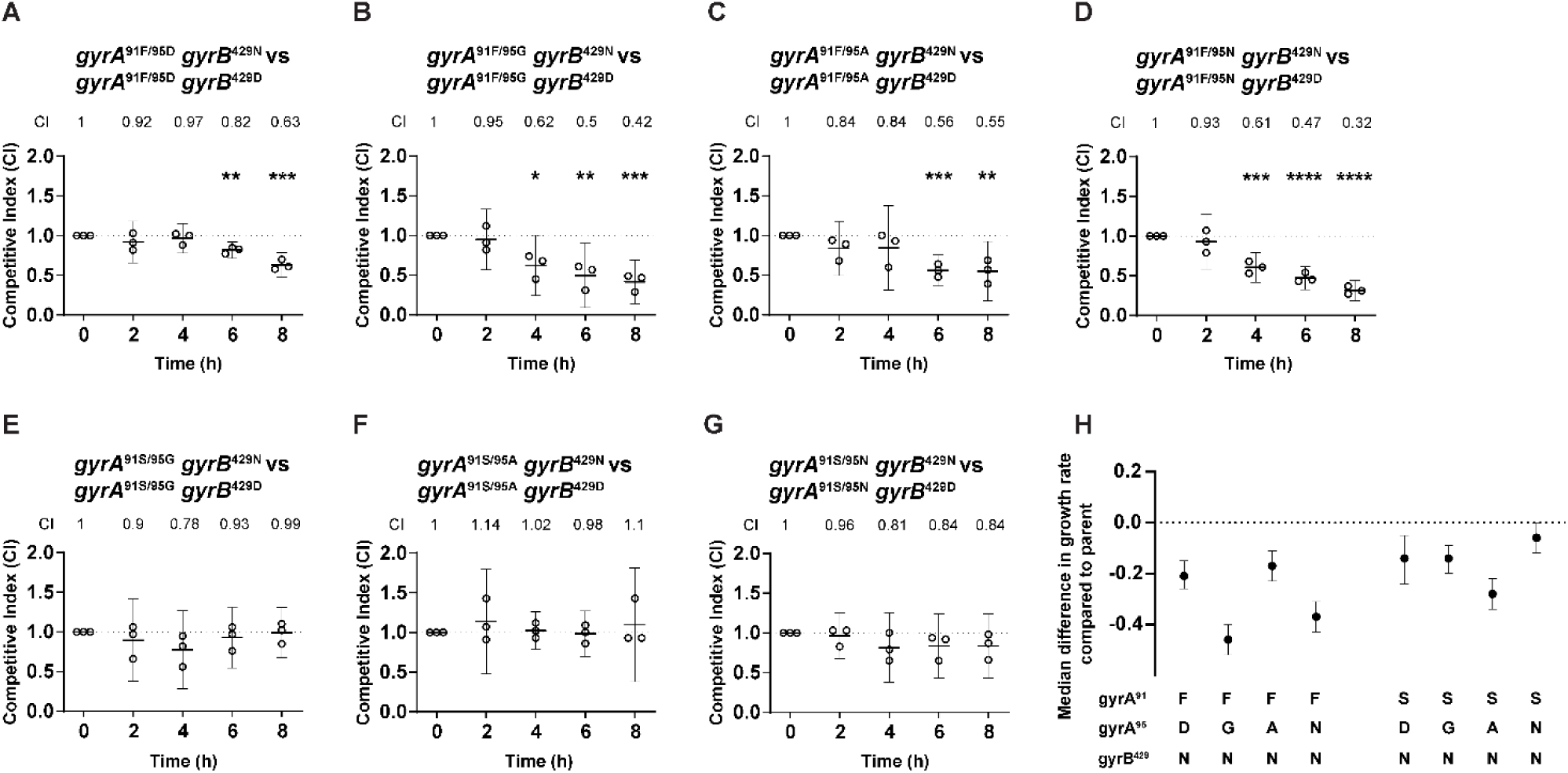
Relative fitness of each *gyrB*^D429N^ mutant relative to its GCGS0481 *gyrA*-matched parent strain. Y-axes show the competitive index (CI) (panels A-G) or fitness difference (panel H) of each *gyrB*^D429N^ mutant relative to the parent. In pairwise assays (panel A to G), the *gyrB*^D429N^ mutant carried a kanamycin marker and was cocultured with its unmarked parental strain; Shown: mean with 95% confidence interval, with statistically significant differences in competitive indices compared to time 0 analyzed using an un-paired Student’s *t-test*, indicated *p ≤ 0.05, **p ≤ 0.01, and ***p ≤ 0.001. n = 3, representative of three independent experiments performed in absence of any antibiotic pressure. (A) *gyrA*^91F/95D^ *gyrB*^429N^: p = 0.21, 0.05, 0.96, 0.02, respectively for 2, 4, 6, and 8 hours. (B) *gyrA*^91F/95G^ *gyrB*^429N^: p = 0.21, 0.15, 0.04, 0.008, respectively for 2, 4, 6, and 8 hours. (C) *gyrA*^91F/95A^ *gyrB*^429N^: p = 0.45, 0.57, 0.02, 0.04, respectively for 2, 4, 6, and 8 hours. (D) *gyrA*^91F/95N^ *gyrB*^429N^: p = 0.18, 0.03, 0.003, 0.0005, respectively for 2, 4, 6, and 8 hours. (E) *gyrA*^91S/95G^ *gyrB*^429N^: p = 0.68, 0.61, 0.4, 0.13, respectively for 2, 4, 6, and 8 hours. (F) *gyrA*^91S/95A^ *gyrB*^429N^: p = 0.15, 0.07, 0.18, 0.24, respectively for 2, 4, 6, and 8 hours. (G) *gyrA*^91S/95N^ *gyrB*^429N^: p = 0.33, 0.11, 0.13, 0.06, respectively for 2, 4, 6, and 8 hours. (H) Estimated difference in exponential-phase growth rate between each strain and its corresponding parent strain, calculated from pooled competition experiments via amplicon sequencing of barcodes; Shown: median and 95% credible interval (CrI) estimate for difference in growth rate, estimated from 2 biological replicates of all barcoded strains grown together (4 barcodes per strain per replicate; timepoints at 0, 2, and 4 hours of coculture).

We next quantified the fitness effects of each *gyrA* allele in absence of the zoliflodacin resistance mutation *gyrB*^D429N^. In pairwise competition with the ciprofloxacin susceptible *gyrA*^91S/95D^ strain, the *gyrA*^S91F^ mutation conferred a modest but measurable fitness disadvantage (**Supplementary Figure 5B**). This fitness disadvantage persisted when the gyrA^S91F^ mutation was combined with substitutions at codon 95 (G, A or N), with the fitness cost magnitude varying somewhat by allele (**Supplementary Figure 5C-E**). In contrast, substitutions at codon 95 in the ciprofloxacin susceptible *gyrA*^91S^ background did not confer a detectable fitness disadvantage in pairwise assays (**Supplementary Figure 5F-H**). In each case, pairwise competition results were consistent with the reciprocal competition in which the kanamycin marker was carried by the opposite strain (**Supplementary Figure 6A-G**). However, when all strains competed in a single pool, fitness estimates for most *gyrA* alleles by barcode sequencing were either neutral (*gyrA*^91F/95D^, *gyrA*^91F/95G^, and *gyrA*^91S/95A^) or low-level fitness reductions (*gyrA*^91F/95A^, *gyrA*^91F/95N^, *gyrA*^91S/95G^, and *gyrA*^91S/95N^) relative to the wild-type strain (**Supplementary Figure 5H**).

### *gyrA*^95^ mutations incur distinct fitness costs in strains with both *gyrB*^D429N^ and *gyrA^S91F^*

Because the use of ciprofloxacin and zoliflodacin may select for strains resistant to both drugs, we next tested whether substitutions at *gyrA*^95^ modulate the fitness costs conferred by *gyrA*^91F^ and *gyrB*^D429N^ in combination. We performed pairwise competition assays between isogenic strains differing only at *gyrA*^95^ in a fixed *gyrA*^91F^ background. *gyrA*^95^ mutants showed no significant fitness defect in pairwise fitness assays in the absence of *gyrB*^D429N^ (**Figure 3A-C**). However, in the presence of *gyrB*^D429N^, *gyrA*^D95G^ and *gyrA*^D95N^ substitutions conferred reduced fitness compared to *gyrA*^91F/95D^ (**Figure 3D,F**). *gyrA*^D95A^ also conferred a modest fitness cost compared to *gyrA*^91F/95D^, although the defect was observed only during stationary phase (**Figure 3E**), and the fitness disadvantage was less severe than that of *gyrA*^D95G^ and *gyrA*^D95N^. As above, all pairwise competition results held when the reciprocal competition was performed in which the kanamycin marker was carried by the opposite strain (**Supplementary Figure 7A-F**).

**Figure 3:**
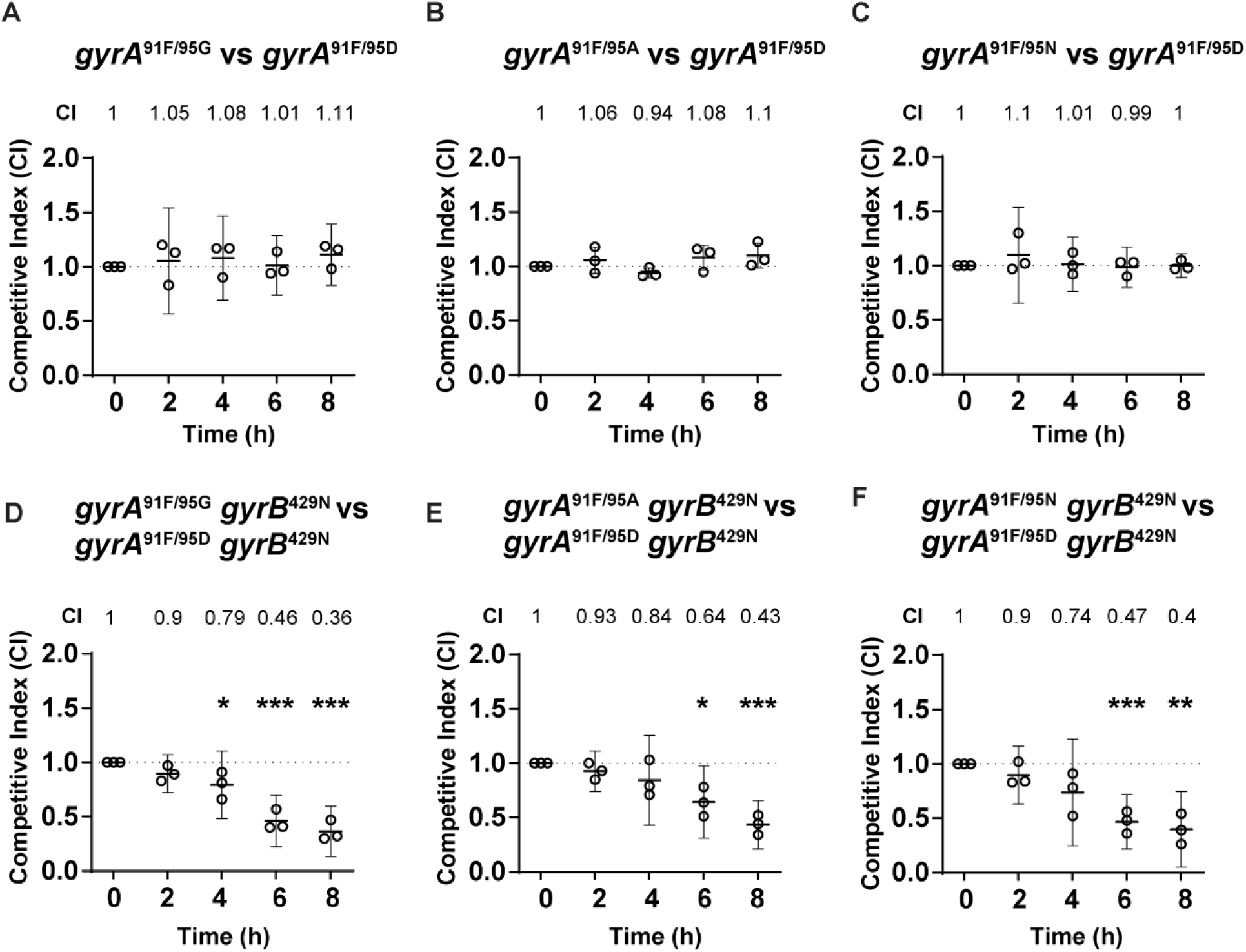
Relative fitness of GCGS0481 *gyrA*^S91F^ strains with various *gyrA*^95^ alleles in the presence or absence of *gyrB*^D429N^ mutation. Y-axes show the competitive index (CI) of each *gyrA*^95^ genotype relative to GCGS0481 *gyrA*^S91F/95D^ in the absence of *gyrB*^D429N^ (A-C) or in the presence of *gyrB*^D429N^ (D-F). (A) *gyrA*^91F/95G^: p = 0.45, 0.28, 0.48, 0.13, respectively for 2, 4, 6, and 8 hours. (B) *gyrA*^91F/95A^: p = 0.12, 0.78, 0.08, 0.06, respectively for 2, 4, 6, and 8 hours. (C) *gyrA*^91F/95N^: p = 0.15, 0.18, 0.21, 0.09, respectively for 2, 4, 6, and 8 hours. (D) *gyrA*^91F/95G^ *gyrB*^429N^: p = 0.32, 0.12, 0.003, 0.001, respectively for 2, 4, 6, and 8 hours. (E) *gyrA*^91F/95A^ *gyrB*^429N^: p = 0.52, 0.3, 0.03, 0.003, respectively for 2, 4, 6, and 8 hours. (F) *gyrA*^91F/95N^ *gyrB*^429N^: p = 0.69, 0.44, 0.02, 0.01, respectively for 2, 4, 6, and 8 hours. n = 3, representative of three independent experiments per-formed in absence of any antibiotic pressure. Shown mean with 95% confidence interval, with statistically significant differences in competitive indices compared to time 0 analyzed using an unpaired Student’s *t-test*,*p ≤ 0.05 and **p ≤ 0.01.

## Discussion

Two new antibiotics, zoliflodacin and gepotidacin, are topoisomerase inhibitors that can be effective against multidrug-resistant uncomplicated urogenital *N. gonorrhoeae* infections^9, 28^. How best to deploy these antibiotics and the expected duration of their clinically useful lifespans depend on the extent to which resistance to each drug is independent of each other and of existing resistance pathways^29, 30^. Despite their distinct mechanisms of action, there is potential for cross-resistance between zoliflodacin and ciprofloxacin^13^ and between zoliflodacin and gepotidacin^25^. Here, we examined epistatic interactions between resistance-associated *gyrA* and *gyrB* variants by systematically reconstructing common ciprofloxacin resistance-associated *gyrA* variants with and without the zoliflodacin resistance mutation *gyrB*^D429N^. We found that these resistance mutations contributed to cross-resistance between ciprofloxacin and zoliflodacin and further showed that the phenotypic effects of *gyrB*^D429N^ were not uniform across backgrounds.

While *gyrB*^D429N^ imposed a fitness cost in all backgrounds tested, the severity of this cost varied with *gyrA* genotype. This variation represented true nonadditive genetic interaction between *gyrB*^D429N^ and *gyrA* alleles, as the cost conferred by *gyrB*^D429N^ varied among strains even after accounting for *gyrA* fitness effects. Among strains carrying the *gyrA*^S91F^ variant, the combination of *gyrA*^D95G^ or *gyrA*^D95N^ with *gyrB*^D429N^ resulted in pronounced fitness costs, whereas the *gyrA*^91F/95A^ background largely mitigated the cost of *gyrB*^D429N^ *in vitro*. These results are consistent with the findings that, in a panel of 9 phylogenetically diverse ciprofloxacin-resistant clinical isolates, *gyrB*^D429N^ conferred more severe fitness costs in strains with the *gyrA*^S91F/95G^ allele than in strains carrying *gyrA*^S91F/95A^ ^25^.

As new drugs such as zoliflodacin are likely to be needed most in areas where ceftriaxone resistance levels are highest, and these areas also have high levels of ciprofloxacin resistance, it is likely that zoliflodacin resistance will emerge from bacterial populations in which ciprofloxacin resistance is common. Our results suggest that not all ciprofloxacin-resistant strains are equally likely to give rise to successful zoliflodacin resistant lineages and that strains with *gyrA*^91F/95A^ may represent the highest risk for development of zoliflodacin resistance via the *gyrB*^D429N^ mutation. Recent phylodynamic analyses have shown that *gyrA*^91F/95A^ lineages expanded more rapidly than *gyrA*^91F/95G^ following the withdrawal of fluoroquinolones from first-line therapy^31^. Together, these findings suggest that the *gyrA*^S91F/95A^ genotype represents both a less costly pathway to ciprofloxacin resistance and a genetic background in which zoliflodacin resistance can arise with comparatively lower fitness costs.

Amplicon sequencing-based approaches can complement standard two-strain competition assays. Amplicon sequencing avoids noise introduced by plating efficiency, allows multiplexing of strains without needing additional markers for subtraction plating, and provides a straightforward path to scaling up biological replicate measurements by including multiple barcodes per strain of interest. The lower complexity of pairwise competition mixtures permits extended co-culture experiments that included periods of culture adaptation, exponential growth, and stationary phase. As a result, subtle late-phase or cumulative effects may be apparent only in pairwise competitions. The complementary nature of these assays can enable a more nuanced dissection of how resistance mutations may impact fitness across physiological states. Concordance between the two approaches indicates fitness effects that manifest during exponential growth, while discrepancies can generate hypotheses for testing fitness in additional growth phase-specific experiments. The partial discordance between the two assays likely also reflects the limited sensitivity of pairwise competition to detect small exponential-phase growth defects. Taken together, these results support a model in which the fitness of zoliflodacin resistant strains is modulated by interactions between *gyrB*^D429N^ and *gyrA* alleles. The most pronounced fitness costs appear driven by interactions between *gyrB*^D429N^ and specific *gyrA* resistance alleles, particularly *gyrA*^91F/95G^ and *gyrA*^91F/95N^. This conclusion is further supported by monoculture growth analyses, in which only the *gyrA*^91F/95G^ *gyrB*^D429N^ and *gyrA*^91F/95N^ *gyrB*^D429N^ strains displayed measurable growth defects (**Supplementary Figures 1 and 2**), and by the concordance between amplicon sequencing and pairwise competition results for these strains.

This study has limitations. All experiments were conducted *in vitro*, and *in vivo* conditions may alter fitness phenotypes and resistance evolutionary trajectories. Experiments were conducted in a single genetic background (GCGS0481 with *parC*^S87R^), and we examined only a subset of clinically observed resistance determinants, focusing on *gyrA* codons 91 and 95 and a single zoliflodacin-resistance mutation, *gyrB*^D429N^. As a result, the epistatic patterns and cross-resistance effects we observed may not generalize to other common gonococcal lineages or to alternative *gyrB* and *parC* alleles, and our short-term *in vitro* competition assays may not fully capture longer-term evolutionary dynamics or phenotypes such as transmission efficiency, persistence, or interactions with other resistance mechanisms.

As zoliflodacin and gepotidacin move into routine clinical use, it will be critical to monitor in real time how resistance emerges in the context of pre-existing topoisomerase variation. Genomic surveillance that jointly tracks *gyrA*, *gyrB*, and *parC* genotypes alongside clinical outcomes will be essential to detect early signals of cross-resistance and to identify strain backgrounds in which zoliflodacin and gepotidacin resistance mutations are most likely to arise and spread. More broadly, the concern that shared resistance pathways can link legacy drugs such as ciprofloxacin to new agents suggests that future drug development for gonorrhea should prioritize compounds that target distinct aspects of the bacterial cellular machinery, with minimal overlap in resistance determinants. Designing and advancing candidates with non-redundant targets and low potential for cross-resistance may be key to prolonging the useful therapeutic lifespan of new antibiotics in gonococcal populations that are already highly enriched for AMR.

## Methods

### *N. gonorrhoeae* culture conditions

*N. gonorrhoeae* was cultured on GCB agar (Difco) supplemented with Kellogg’s supplement (GCB-K) at 37°C with 5% CO_2_^32^. Growth curve and relative fitness assessments were conducted in liquid GCP medium containing 15 g/L proteose peptone 3 (Thermo Fisher), 1 g/L soluble starch, 1 g/L KH_2_PO_4_, 4 g/L K_2_HPO_4_, and 5 g/L NaCl (Sigma-Aldrich) with Kellogg’s supplement^33^ with agitation at 37°C with 5% CO_2_.

### Generation of isogenic *N. gonorrhoeae gyrA* and *gyrB* mutant strains

Strains, plasmids and primers used in this study are listed in **Supplementary Table 1**. Isogenic *N. gonorrhoeae* strains were generated in a ciprofloxacin-resistant clinical isolate, GCGS0481, which natively carries *gyrA*^91F/95G^ and *parC*^87R^ alleles. A *gyrA*^91S/95D^ derivative of GCGS0481 was constructed with a chloramphenicol resistance (CM^R^) marker as described^31^. The GCGS0481 *gyrA*^91S/95D^ strain harbors the *gyrA* allele from the susceptible laboratory strain FA19, which also contains the *gyrA*^M250I^ variant. This variant has been reported in other *N. gonorrhoeae* genomes^34^ and does not contribute to fluoroquinolone-resistance or impact *in vitro* fitness^31^. Alleles of *gyrA* with resistance-associated substitutions at amino acid positions 91 and 95 were amplified using primers AM_5 (F) and AM_6 (R) from the genomic DNA of clinical *N. gonorrhoeae* isolates NY0464 (*gyrA*^91S/95G^, European Nucleotide Archive or ENA accession: ERR2631942), EEE002 (*gyrA*^91S/95A^, NCBI Sequence Read Archive or SRA accession: SRR16683648), NZ2015-82 (*gyrA*^91S/95N^, SRA accession: SRR5827182), GCGS0709 (*gyrA*^91F/95D^, ENA accession: ERR1067871), GCGS0481(*gyrA*^91F/95G^, ENA accession: ERR855135), NY0842 (*gyrA*^91F/95A^, ENA accession: ERR2631880) and 157M (*gyrA*^91F/95N^, SRA accession: SRR16683705). Each allele was introduced into the chloramphenicol marked GCGS0481 *gyrA*^91S/95D^ strain by electroporation^31^, and individual colonies were selected on GCB-K plates supplemented with 0.25 µg/mL ciprofloxacin (*gyrA*^91S/95G^, *gyrA*^91S/95A^, *gyrA*^91S/95N^) or 2 µg/mL ciprofloxacin (*gyrA*^91F/95D^, *gyrA*^91F/95G^, *gyrA*^91F/95A^, *gyrA*^91F/95N^). Transformants were screened by Sanger sequencing to ensure that the recombined region did not affect amino acid 250, the nearest variable site in *gyrA*. Thus, all strains in the isogenic panel contained the *gyrA*^M250I^ variant and are comparable.

The mutant *gyrB*^D429N^ allele was amplified using primers AM_1 (F) and AM_2 (R) from the genomic DNA of an experimentally evolved *N. gonorrhoeae* GCGS0481 strain that acquired the *gyrB*^D429N^ mutation under ciprofloxacin pressure^13^ and introduced into the isogenic GCGS0481 strains by electroporation^25, 31^. Individual colonies were selected on GCB-K plates from within the zone of inhibition created by a dried droplet of 4 μg/mL zoliflodacin. For all the transformations performed, transformations without DNA were used as negative controls. Transformants were verified by Sanger sequencing.

### Antibiotic susceptibility testing

Antibiotic susceptibility testing was performed on GCB-K agar via MIC test strips (Bio-Merieux and Liofilchem) for ciprofloxacin or agar dilution for zoliflodacin, gepotidacin, and nalidixic acid. All MIC results reported are the mean of three independent experiments.

### Growth curve assays

To measure the growth of the isogenic GCGS0481 strains, overnight cultures on GCB-K plates were diluted to an optical density (OD) of 0.1 (600nm) and grown in antibiotic-free GCP media with Kellogg’s supplement for 8 hours. OD_600_ of each strain was measured at 2, 4, 6 and 8 hours using a spectrophotometer (Thermo Scientific). At each timepoint, cultures were also serially diluted and plated on GCB-K agar plates to measure colony forming units (CFUs) as a proxy for viable bacteria.

### Assessment of relative fitness

#### Pairwise fitness assays

The pairwise competition assays were performed as described^25^. Kanamycin resistance was introduced into each strain in the panel of GCGS0481 isogenic strains with pDR53, which integrates an *aphA3* marker onto the chromosome between *lctP* and *aspC*^31^. Transformants were selected on GCB-K agar supplemented with 70 µg/ml kanamycin. Paired strains (one kanamycin-sensitive and one kanamycin-resistant strain) were mixed at a ratio of 1:1 by OD_600_, diluted to OD_600_ ∼0.1, and co-cultured in 5-10 mL of antibiotic-free GCP media with Kellogg’s supplement for 8 hours. At each timepoint, cultures were serially diluted and plated on both GCB-K agar and GCB-K agar supplemented with 70 µg/ml kanamycin. Plates were incubated overnight at 37°C with 5% CO_2_. Colonies on each plate were quantified, and the competitive index was calculated at each timepoint as (*R_t_* / *S_t_*) / (*R_0_* / *S_0_*), where *R_t_*and *S_t_* are the proportions of kanamycin-resistant and kanamycin-sensitive strains, respectively, at time *t* and *R_0_* and *S_0_* are the proportions of kanamycin-resistant and kanamycin-sensitive strains at time 0. A competitive index value of 1 indicates equal fitness between strains, >1 indicates the kanamycin-resistant mutant is more fit than the parental strain, and <1 indicates the mutant is less fit. Statistical analysis of competitive index measurements was performed using an unpaired two-sided Student’s *t-test*. All the pairwise competition assays were performed with reciprocal kanamycin-marking to ensure that the kanamycin marker did not impact reported fitness.

#### Fitness comparison using amplicon sequencing

##### Creation of the barcoding plasmid

The plasmid pBC1 was constructed to introduce a random 20-mer barcode with a kanamycin marker into the neutral site on the gonococcal chromosome between *aspC* and *lctP*. pBC1 was constructed by cloning 20-mer barcodes into the PacI/XbaI site of pDR53^25, 31^. Briefly, an insert was created by amplifying the primer DR412 (containing a 20-mer of random nucleotides flanked by homology to the pDR53 multiple cloning site) with a short complementary primer DR413 by PCR with Taq polymerase (NEB) for 10 cycles (95°C for 22 seconds, 50°C for 37 seconds, 68°C for 15 seconds). The resulting product and the pDR53 plasmid were digested with XbaI and PacI (NEB), ligated using T4 ligase (NEB), and transformed into chemically competent DH5𝘢. To maintain diversity of barcodes, plasmids were purified from a pool of ∼120,000 independent transformants.

##### Barcoding of isogenic strain panel

The barcoding plasmid was transformed into each GCGS0481-derived strain in the isogenic panel via electroporation^31^. Transformants were selected on GCB-K and 70 µg/mL kanamycin. Barcode sequences were amplified by PCR with Taq polymerase and primers P6 and P7, and Sanger sequenced using primer P7. Four clones with unique barcode sequences were chosen and pooled for each strain background.

##### Amplicon sequencing competition assay

For each strain to be included in the competition assay, a pool of four individual bar-coded derivatives was grown on GCB-K plates at 37°C with 5% CO_2_ for 16-18 hours and suspended in liquid GCP supplemented with Kellogg’s to OD_600_ 0.3 in ∼2 mL. Suspensions were incubated with shaking at 37°C with 5% CO_2_ for ∼1.5 hours to allow cultures to return to early exponential phase. Strains with particularly poor overnight growth (at most 2 per experiment) were necessarily inoculated at a lower density and received approximately 1 additional hour of outgrowth. After outgrowth, OD_600_ measurements were repeated, and a pool was made of all strains equalized by OD. The competition pool was diluted to OD_600_ 0.01 in 15 mL of fresh GCP-K and incubated for 8 hours, shaking, at 37°C with 5% CO_2_. Every two hours, an aliquot was taken for barcode abundance measurement and dilutions were plated on GCB-K for CFU estimation. Two independent replicates of the competition experiment were performed.

Aliquots from each time point were pelleted by centrifugation for one minute at 21100×*g* and pellets were frozen at -80°C. To amplify barcodes, genomic DNA was extracted from thawed pellets using the Invitrogen™ PureLink™ Genomic DNA Mini Kit and 20 cycles of PCR were performed using 50 ng of gDNA, Taq polymerase (NEB), and primers P41 and P42 (**Supplementary table 1**). To reduce amplicon jackpotting, PCR was performed in triplicate. Replicate reactions were pooled and products were purified with the QIAquick PCR Purification Kit (Qiagen) and eluted in nuclease-free water. A sequencing library was prepared from the barcode amplicons with the Oxford Nanopore Native Barcoding Kit 24 v. 14 (SQK-NBD114.24) and sequenced on a MinION Mk1B device. Sequencing continued until each sample had greater than 64k reads (allocating 1000 reads per strain barcode if all strains were present at equal abundance). Nanopore reads were basecalled with Dorado v0.8.1 (https://github.com/nanoporetech/dorado) with the super accuracy model. The resulting fastq files were passed through a custom python script (available at https://github.com/gradlab/barcoded_fitness_gyrA-gyrB) to count the instances of each barcode.

##### Calculation of genotype fitness from barcode data

Fitness was calculated as a function of change in barcode abundance over time, with time expressed as generations of growth of the total strain pool (calculated from dilution plating during the competition). Only the 0, 2, and 4-hour timepoints were used, as dilution plating revealed that these were the timepoints during which the culture was undergoing exponential growth. Fitness associated with each barcode was calculated relative to the wild type strain (GCGS0481 *gyrA*^91S/95D^). Briefly, the sequence counts of each barcode at each timepoint for a given experimental replicate were divided by the sum of the sequence counts from the 4 wild type barcodes in the same sample. The variable 𝜋_𝑖,𝑗,𝑘,𝑙_ is this proportion for a strain with *gyrA* genotype *i* and *gyrB* genotype *j* with individual barcode *k* in experimental replicate *l*. Relative fitness was determined with the following regression model using brms with a Gaussian response^35^ :

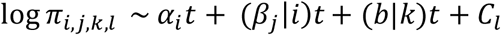

where *t* is time in generations; 𝛼 is the fitness effect of *gyrA* genotype *i*; *β_j_*|*i* is the fitness effect of *gyrB* genotype *j* in a strain carrying *gyrA* genotype *i*; 𝑏|𝑘 is the fitness effect of individual barcode *k* (of the 4 unique barcodes per genotype); and *C_l_*is the error for experimental replicate *l*. Relative fitness, defined as the difference in growth rate between a given genotype and the wild type strain, was estimated as the posterior median of model draws for the fitness effects of the strain’s *gyrA* and *gyrB* genotypes. Fitness of *gyrB*^D429N^ mutants compared to their *gyrB*^429D^ parents were quantified as the posterior median for the *gyrB* term alone. An R file for recreating the analysis is available at https://github.com/gradlab/barcoded_fitness_gyrA-gyrB.

## Acknowledgements

We would like to thank members of the Grad lab for helpful discussions and feedback on this manuscript. This work was supported by NIH R01 AI132606 and R01 AI153521 grants to YHG.

## Author contributions

AM, SGP and YHG conceptualized the study. AM, SOPB, DH, DHFR and BB acquired/analyzed the data. AM, SGP and YHG wrote the original draft, and all the authors reviewed and edited the manuscript.

## Competing interests

YHG has consulted GSK on topics not related to the material in this manuscript.

## Supplementary Files

**Supplementary Table 1:**
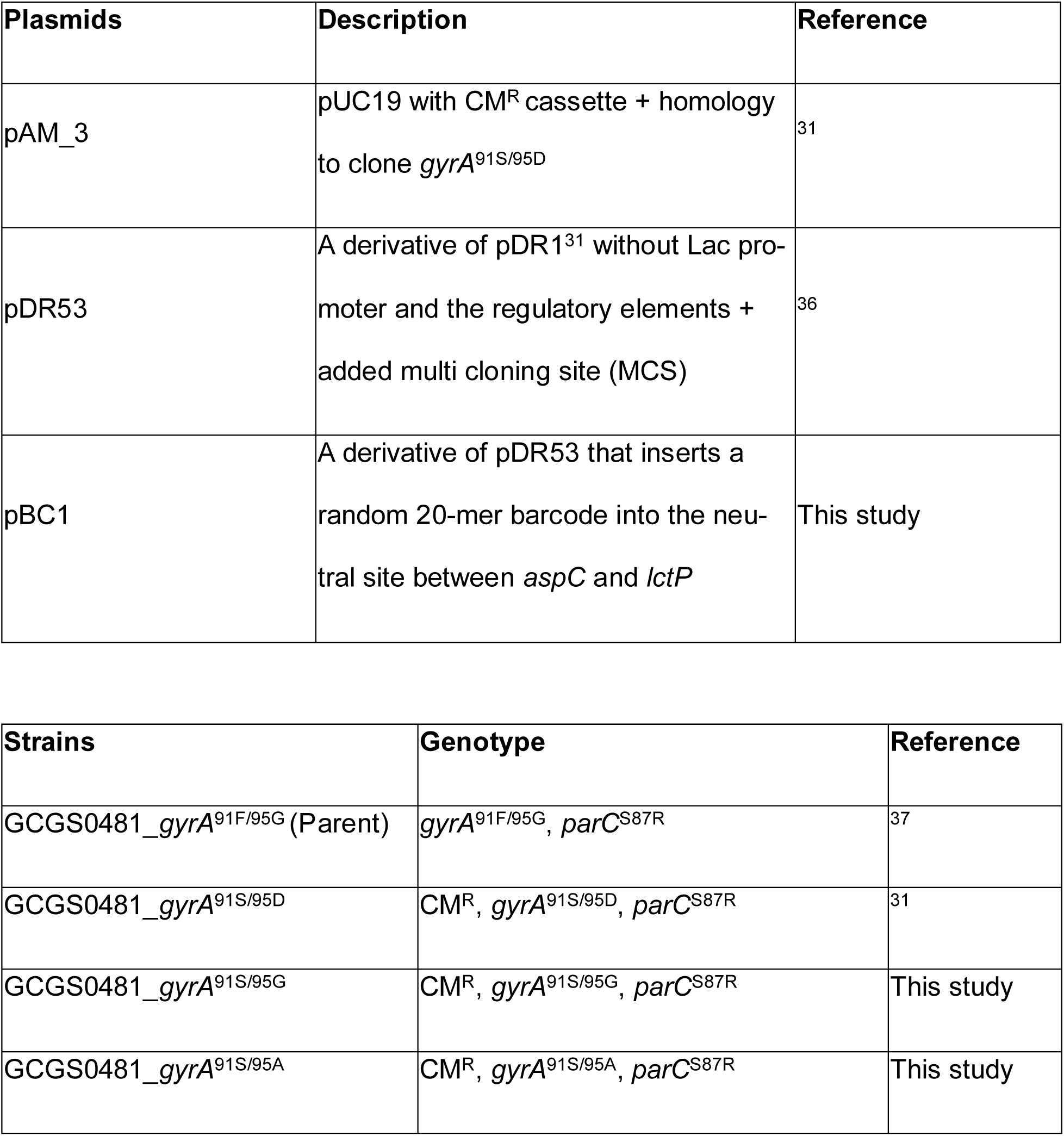
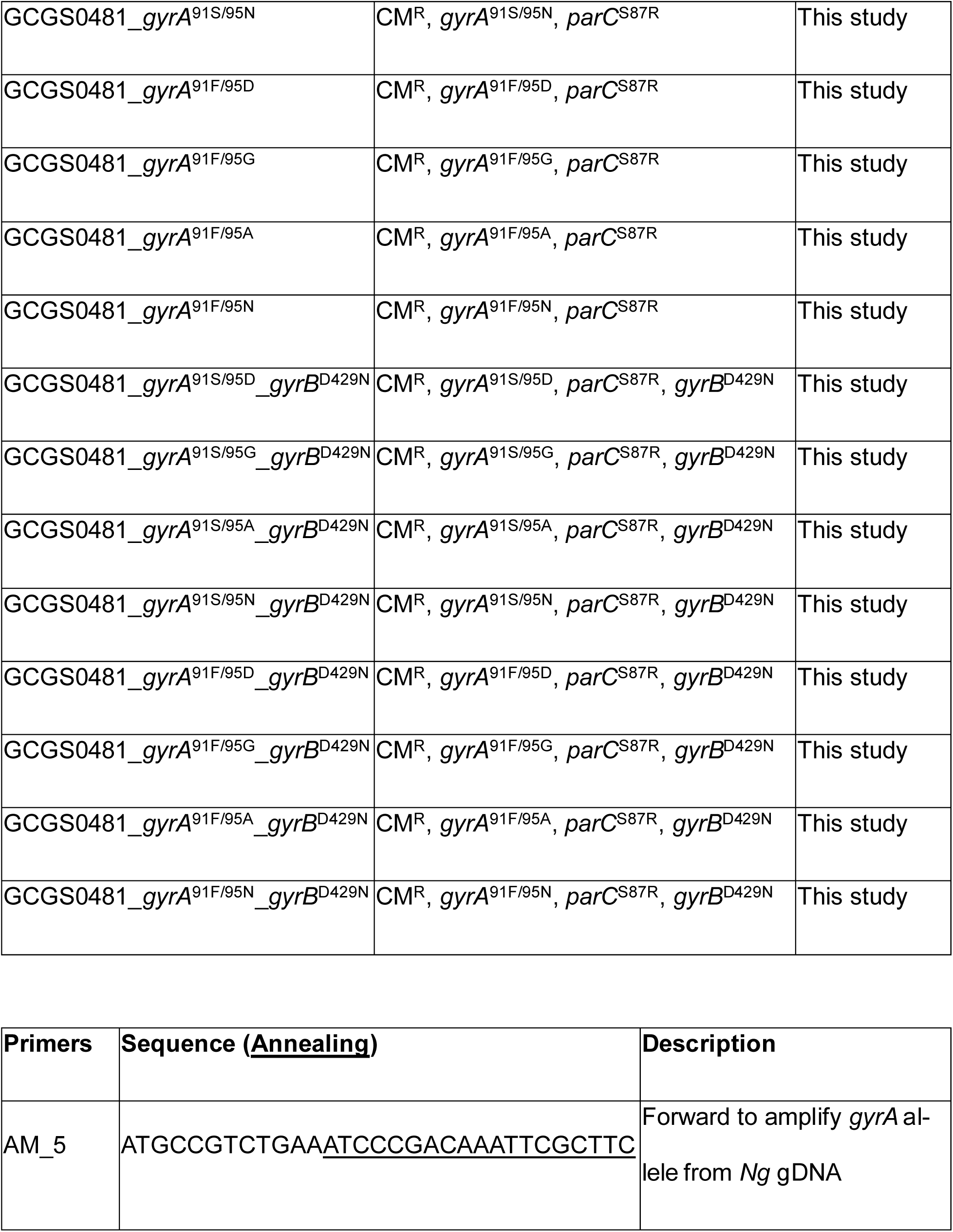
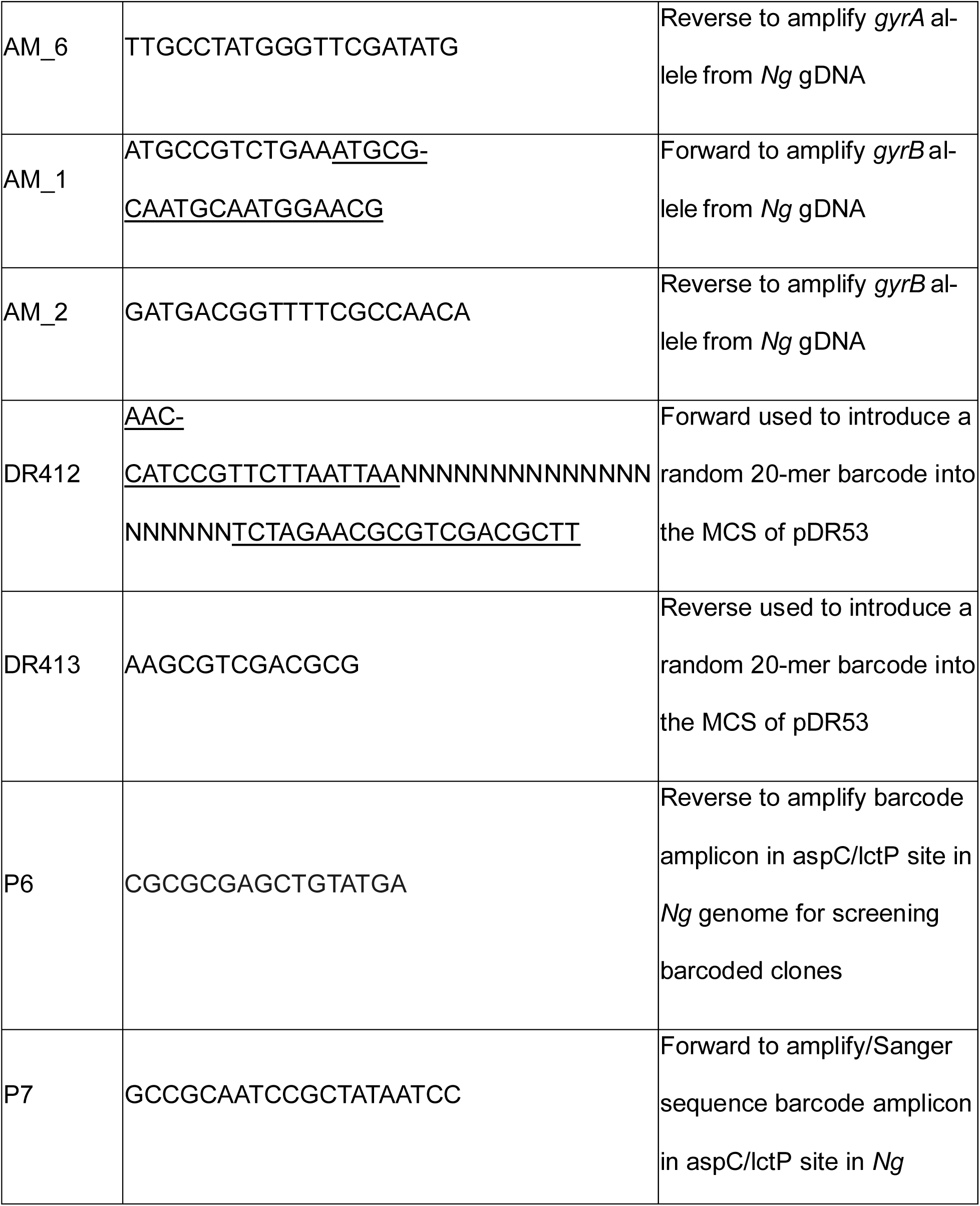
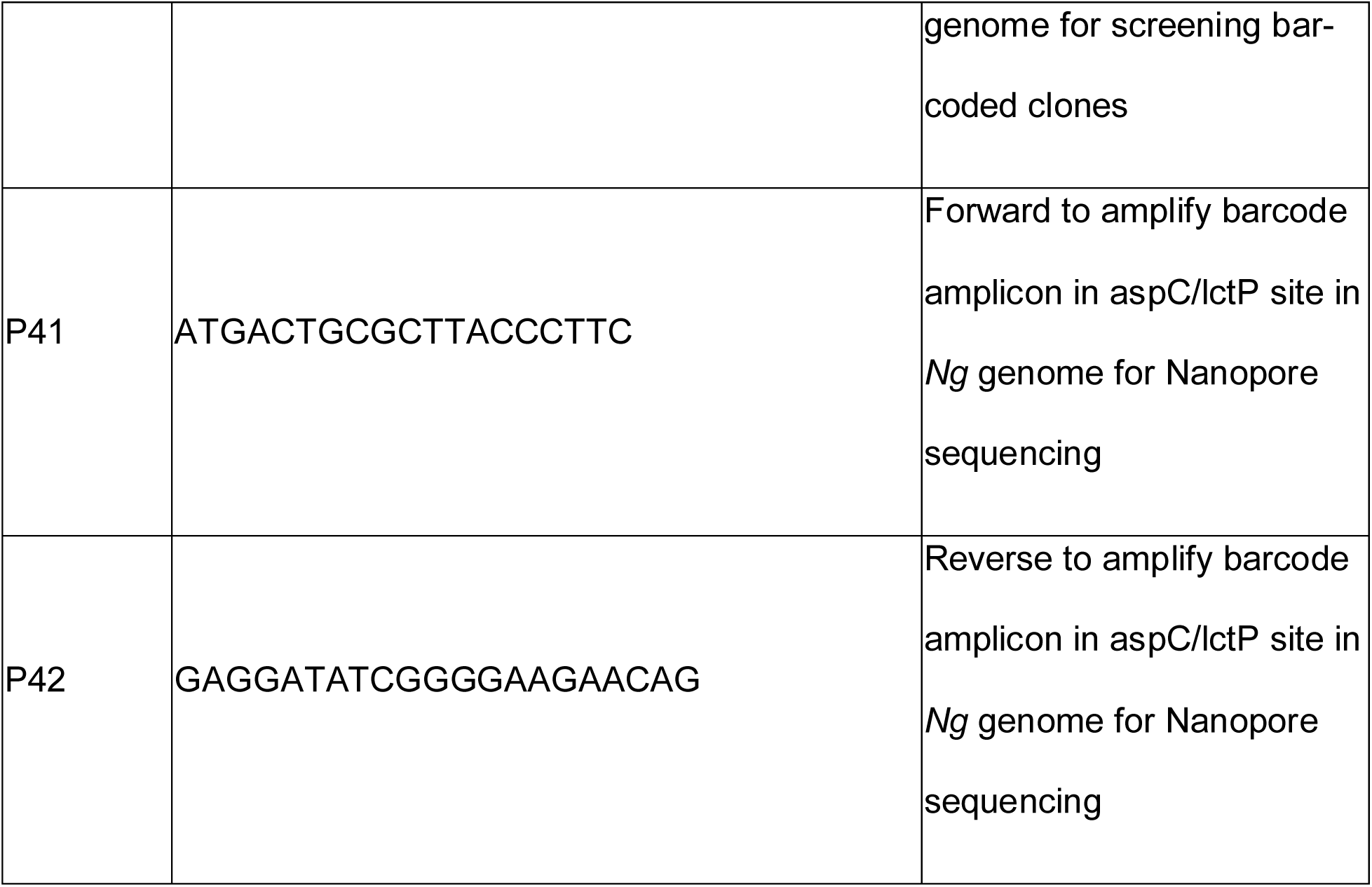
Plasmids, *N. gonorrhoeae* strains and primers used in this study.

**Supplementary Figure 1:**
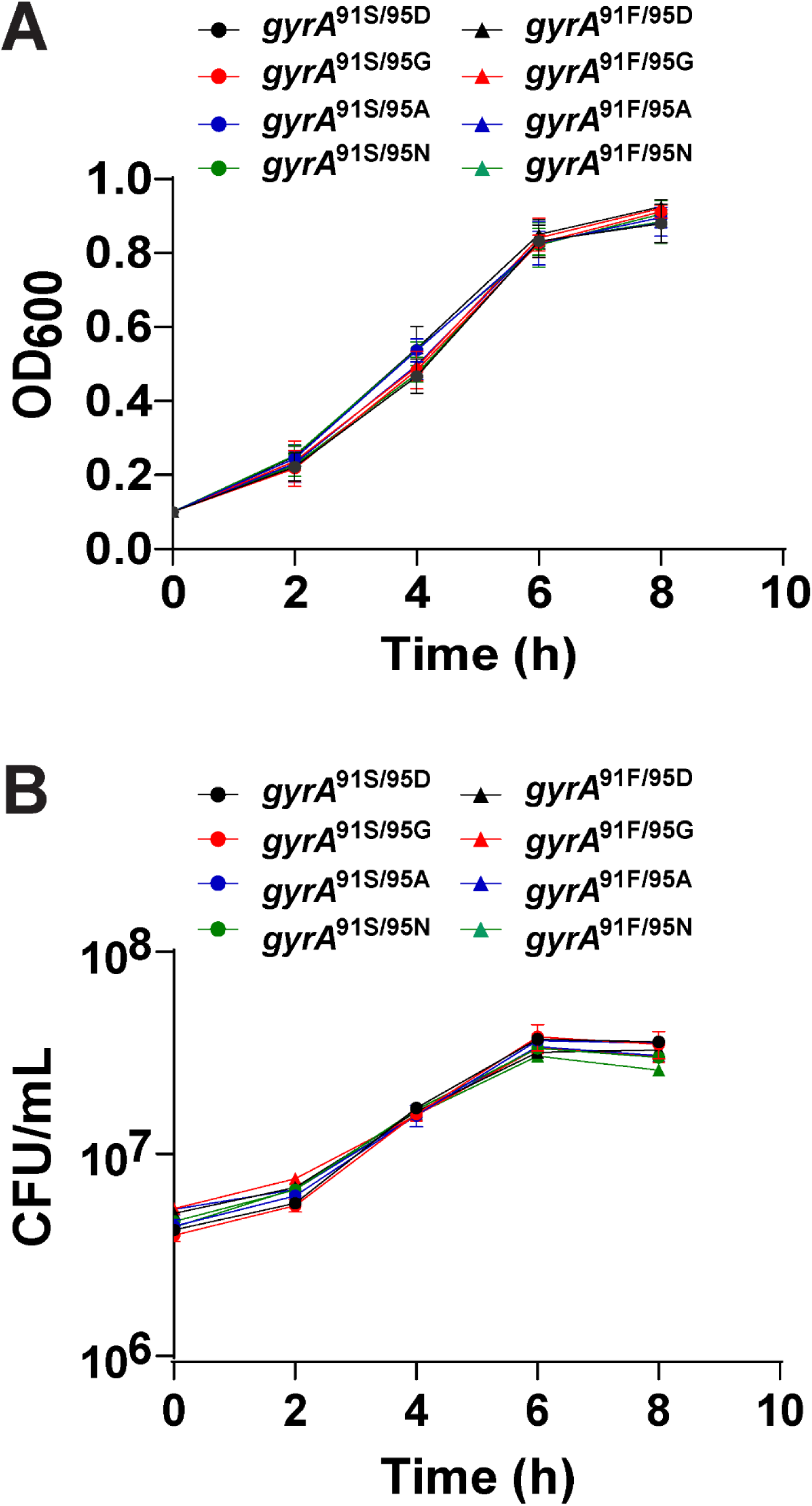
*In vitro* growth kinetics of isogenic strains with different *gyrA* alleles, measured by spectrophotometry. All isogenic strains were cultured separately in liquid GCP media supplemented with Kellogg’s supplement at a starting absorbance reading at 600nm (OD_600_) = 0.1. OD_600_ readings at 0, 2, 4, 6, and 8 hours timepoint are plotted. (A) A_600_ readings at 0, 2, 4, 6, and 8 hours. (B) Dilutions were plated on GCB-K plates at 0, 2, 4, 6, and 8 hours, colonies were counted after overnight growth, and CFUs were calculated. n = 3, representative of three independent experiments. Error bars represent standard deviation of three biological replicates. Statistical significance comparing each strain to GCGS0481 *gyrA*^91S/95D^ was determined by unpaired two-sided Student’s *t*-test (no significant results found).

**Supplementary Figure 2:**
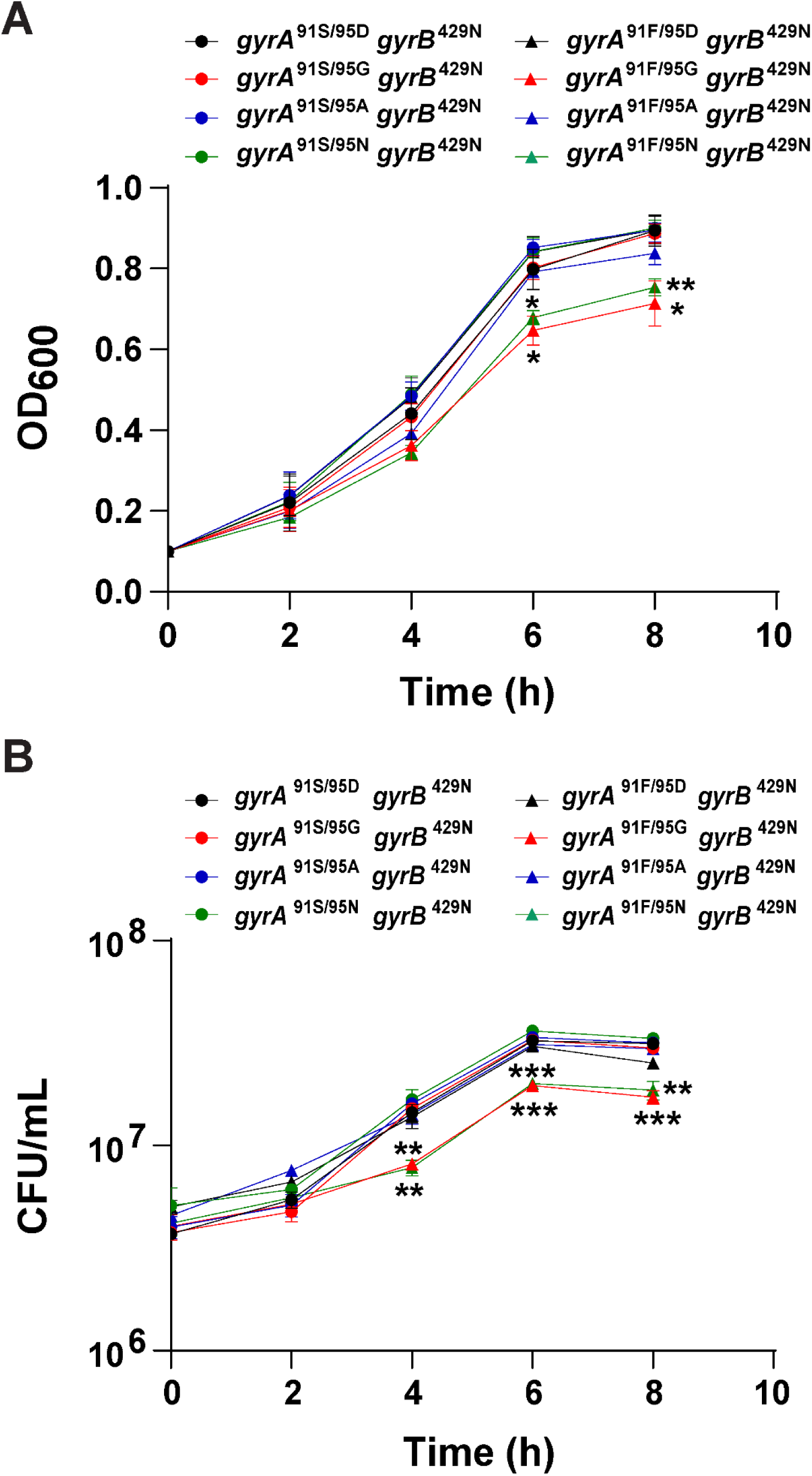
*In vitro* growth kinetics of isogenic strains with *gyrB*^D429N^ in combination with different *gyrA* alleles, measured by spectrophotometry. All isogenic strains were cultured separately in liquid GCP media supplemented with Kellogg’s supplement at a starting absorbance reading at 600nm (OD_600_) = 0.1. OD_600_ readings at 0, 2, 4, 6, and 8 hours timepoint are plotted. (A) A_600_ readings at 0, 2, 4, 6, and 8 hours. (B) Dilutions were plated on GCB-K plates at 0, 2, 4, 6, and 8 hours, colonies were counted after overnight growth, and CFUs were calculated. n = 3, representative of three independent experiments. Error bars represent standard deviation of three biological replicates. Statistical significance comparing each strain to GCGS0481 *gyrA*^91S/95D^ *gyrB*^D429N^ was determined by unpaired two-sided Student’s *t-test* and are indicated *p ≤ 0.05, **p ≤ 0.01, and ***p ≤ 0.001.

**Supplementary Figure 3:**
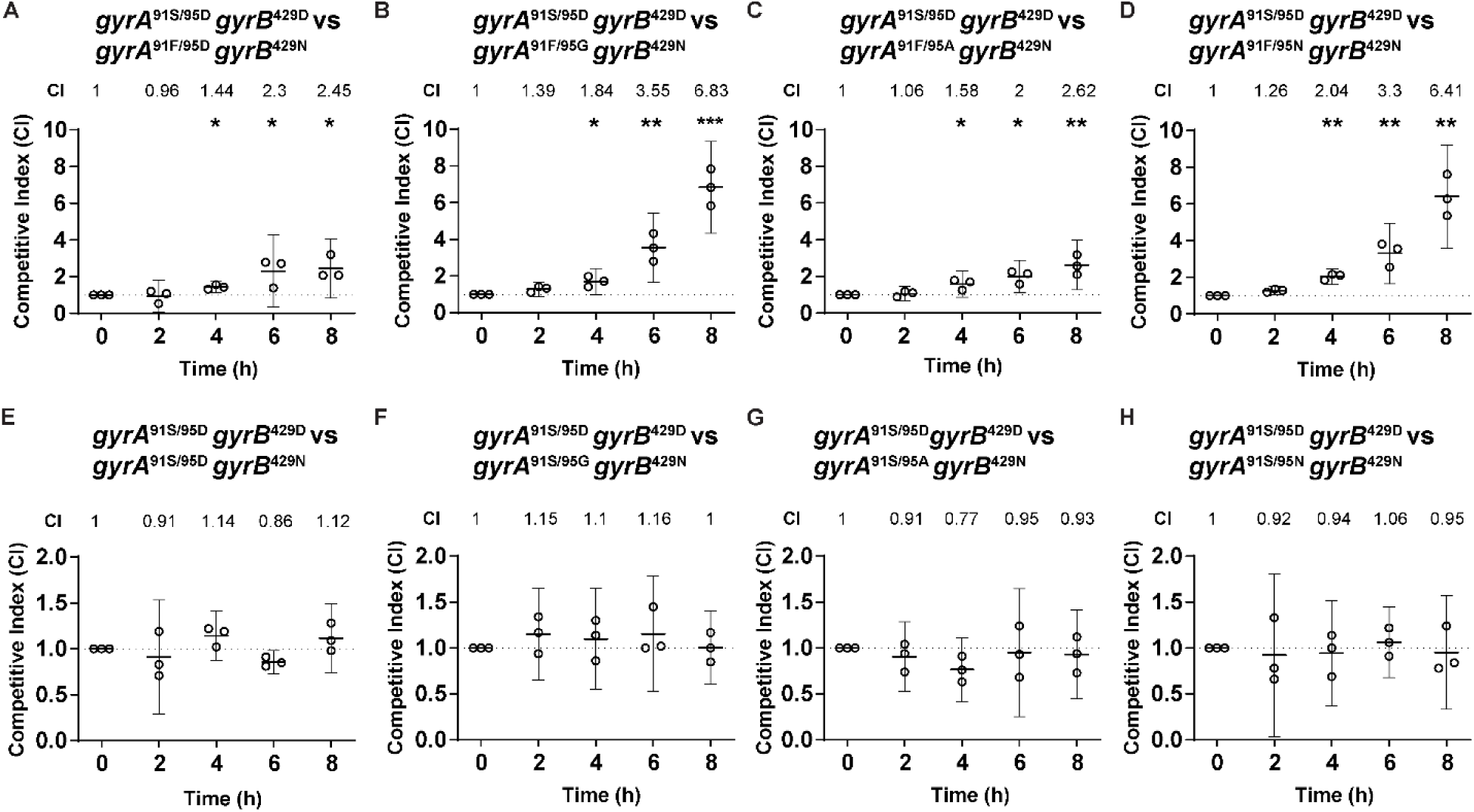
Relative fitness of the *gyrB*^D429N^ mutants with various *gyrA* alleles in the isogenic *gyrA* mutant panel. Y-axes show the competitive index (CI) of the ciprofloxacin-susceptible strain GCGS0481 *gyrA*^91S/95D^ compared to each *gyrB*^D429N^ mutant. The parental strain carried a kanamycin marker and was cocultured with each un-marked *gyrB*^D429N^ mutant strain. (A) *gyrA*^91S/95D^ *gyrB*^429D^ over *gyrA*^91F/95D^ *gyrB*^429N^: p = 0.99, 0.02, 0.04, 0.02, respectively for 2, 4, 6, and 8 hours. (B) *gyrA*^91S/95D^ *gyrB*^429D^ over *gyrA*^91F/95G^ *gyrB*^429N^: p = 0.31, 0.04, 0.006, 0.0006, respectively for 2, 4, 6, and 8 hours. (C) *gyrA*^91S/95D^ *gyrB*^429D^ over *gyrA*^91F/95A^ *gyrB*^429N^: p = 0.97, 0.04, 0.01, 0.008, respectively for 2, 4, 6, and 8 hours. (D) *gyrA*^91S/95D^ *gyrB*^429D^ over *gyrA*^91F/95N^ *gyrB*^429N^: p = 0.09, 0.001, 0.005, 0.001, respectively for 2, 4, 6, and 8 hours. (E) *gyrA*^91S/95D^ *gyrB*^429D^ over *gyrA*^91S/95D^ *gyrB*^429N^: p = 0.92, 0.21, 0.6, 0.29, respectively for 2, 4, 6, and 8 hours. (F) *gyrA*^91S/95D^ *gyrB*^429D^ over *gyrA*^91S/95G^ *gyrB*^429N^: p = 0.32, 0.54, 0.39, 0.98, respectively for 2, 4, 6, and 8 hours. (G) *gyrA*^91S/95D^ *gyrB*^429D^ over *gyrA*^91S/95A^ *gyrB*^429N^: p = 0.51, 0.16, 0.77, 0.64, respectively for 2, 4, 6, and 8 hours. (H) *gyrA*^91S/95D^ *gyrB*^429D^ over *gyrA*^91S/95N^ *gyrB*^429N^: p = 0.81, 0.83, 0.61, 0.88, respectively for 2, 4, 6, and 8 hours. n = 3, representative of three independent experiments performed in absence of any antibiotic pressure. Shown: mean with 95% confidence interval, with statistically significant differences in competitive indices compared to time 0 analyzed using an unpaired Student’s *t-test*, indicated *p ≤ 0.05, **p ≤ 0.01, and ***p ≤ 0.001.

**Supplementary Figure 4:**
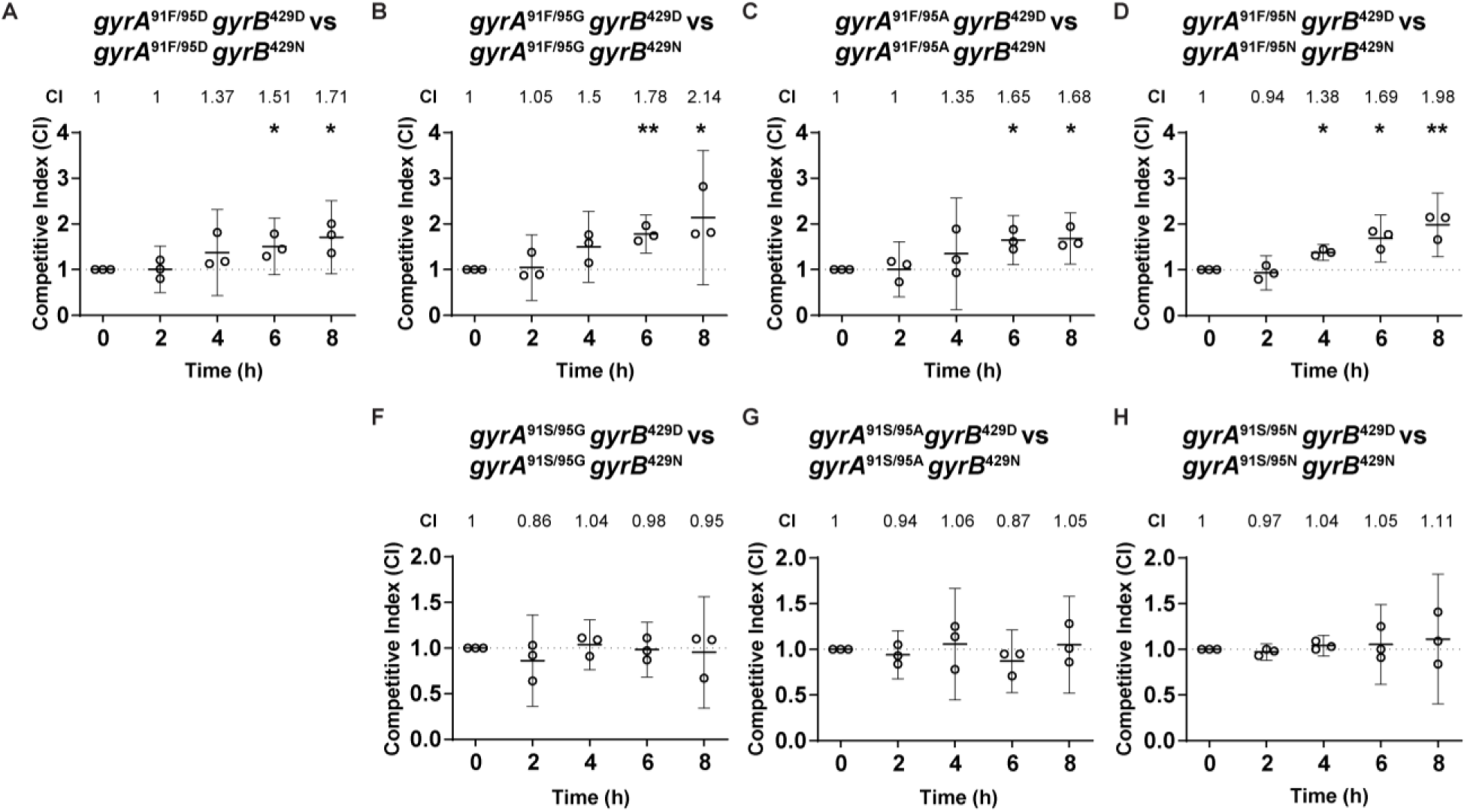
Relative fitness of each *gyrA* strain relative to its own *gyrB*^D429N^ derivative. Y-axes show the competitive index of each (CI) of each parental GCGS0481 relative to *gyrB*^D429N^ mutants. The parental strain carried a kanamycin marker and was cocultured with its unmarked *gyrB*^D429N^ mutant. (A) *gyrA*^91F/95D^ *gyrB*^429D^: p = 0.35, 0.09, 0.02, 0.02, respectively for 2, 4, 6, and 8 hours. (B) *gyrA*^91F/95G^ *gyrB*^429D^: p = 0.99, 0.09, 0.005, 0.04, respectively for 2, 4, 6, and 8 hours. (C) *gyrA*^91F/95A^ *gyrB*^429D^: p = 0.6, 0.42, 0.01, 0.01, respectively for 2, 4, 6, and 8 hours. (D) *gyrA*^91F/95N^ *gyrB*^429D^: p = 0.12, 0.01, 0.01, 0.007, respectively for 2, 4, 6, and 8 hours. (E) *gyrA*^91S/95G^ *gyrB*^429D^: p = 0.44, 0.79, 0.9, 0.81, respectively for 2, 4, 6, and 8 hours. (F) *gyrA*^91S/95A^ *gyrB*^429D^: p = 0.1, 0.15, 0.43, 0.11, respectively for 2, 4, 6, and 8 hours. (G) *gyrA*^91S/95N^ *gyrB*^429D^: p = 0.58, 0.24, 0.36, 0.35, respectively for 2, 4, 6, and 8 hours. n = 3, representative of three independent experiments performed in absence of any antibiotic pressure. Shown: mean with 95% confidence interval. Statistically significant differences in competitive indices compared to time 0 were analyzed using an unpaired Student’s *t-test*, indicated *p ≤ 0.05 and **p ≤ 0.01.

**Supplementary Figure 5:**
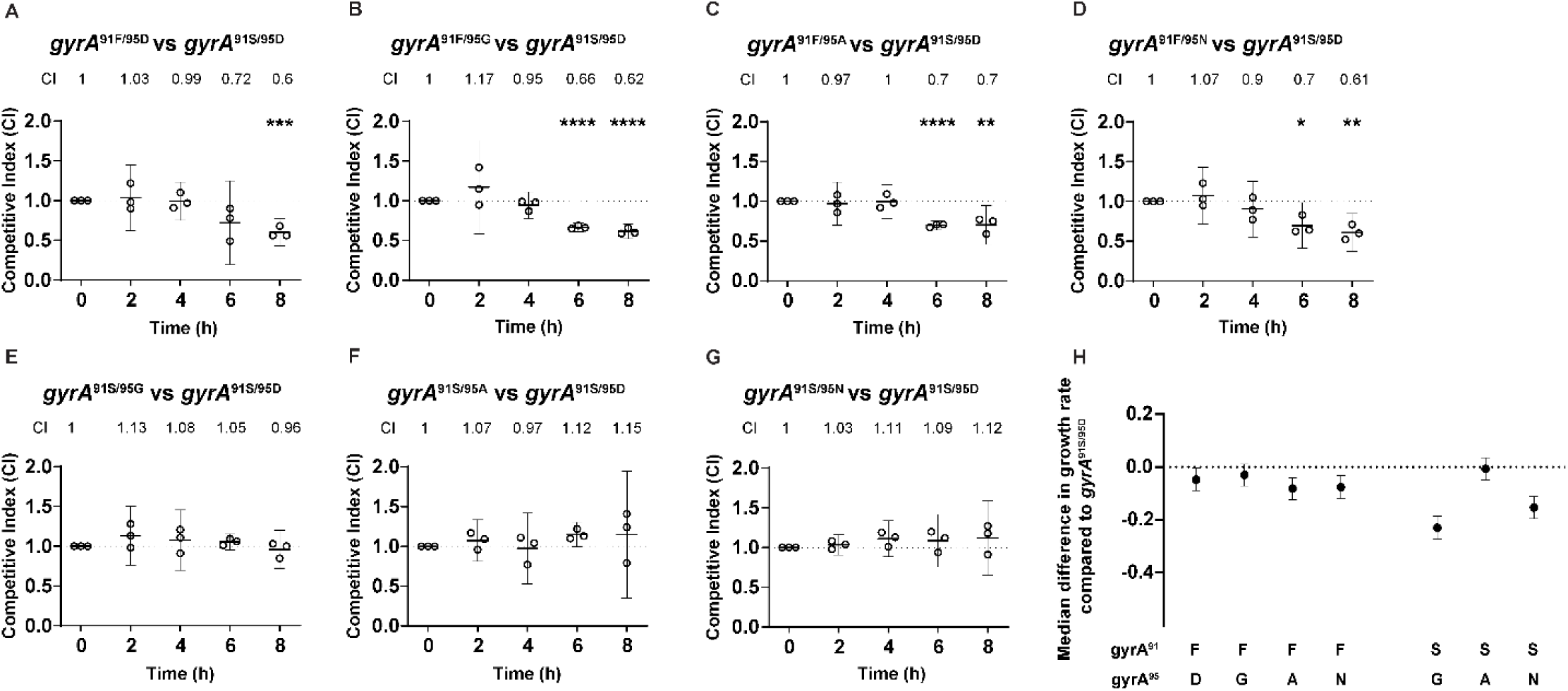
Relative fitness of the each isogenic *gyrA* mutant strain relative to *gyrA*^91S/95D^ in the absence of *gyrB* mutation. Y-axes show the competitive index (CI) (panels A-G) or fitness difference (panel H) of each *gyrA* mutant relative to the ciprofloxacin-susceptible strain *gyrA*^91S/95D^. In pairwise assays (panels A-G), the *gyrA* mutant carried a kanamycin marker and was cocultured with the unmarked *gyrA*^91S/95D^ strain; shown: mean with 95% confidence interval, with statistically significant differences in competitive indices compared to time 0 analyzed using an unpaired Student’s *t-test*, indicated *p ≤ 0.05, **p ≤ 0.01, and ***p ≤ 0.001. n = 3, representative of three independent experiments performed in absence of any antibiotic pressure. (A) *gyrA*^91F/95D^: p = 0.91, 0.25, 0.1, 0.005, respectively for 2, 4, 6, and 8 hours. (B) *gyrA*^91F/95G^: p = 0.24, 0.99, 0.046, 0.03, respectively for 2, 4, 6, and 8 hours. (C) *gyrA*^91F/95A^: p = 0.24, 0.29, 0.0003, 0.005, respectively for 2, 4, 6, and 8 hours. (D) *gyrA*^91F/95N^: p = 0.69, 0.25, 0.01, 0.004, respectively for 2, 4, 6, and 8 hours. (E) *gyrA*^91S/95G^: p = 0.1, 0.21, 0.04, 0.75, respectively for 2, 4, 6, and 8 hours. (F) *gyrA*^91S/95A^: p = 0.48, 0.99, 0.2, 0.47, respectively for 2, 4, 6, and 8 hours. (G) *gyrA*^91S/95N^: p = 0.23, 0.1, 0.05, 0.22, respectively for 2, 4, 6, and 8 hours. (H) Estimated difference in exponential-phase growth rate between *gyrA*^91S/95D^ and each mutant *gyrA* strain, calculated from pooled competition experiments via amplicon sequencing of barcodes; Shown: median and 95% credible interval (CrI) estimate for difference in growth rate, estimated from 2 biological replicates of all barcoded strains grown together (4 barcodes per strain per replicate; timepoints at 0, 2, and 4 hours of coculture).

**Supplementary Figure 6:**
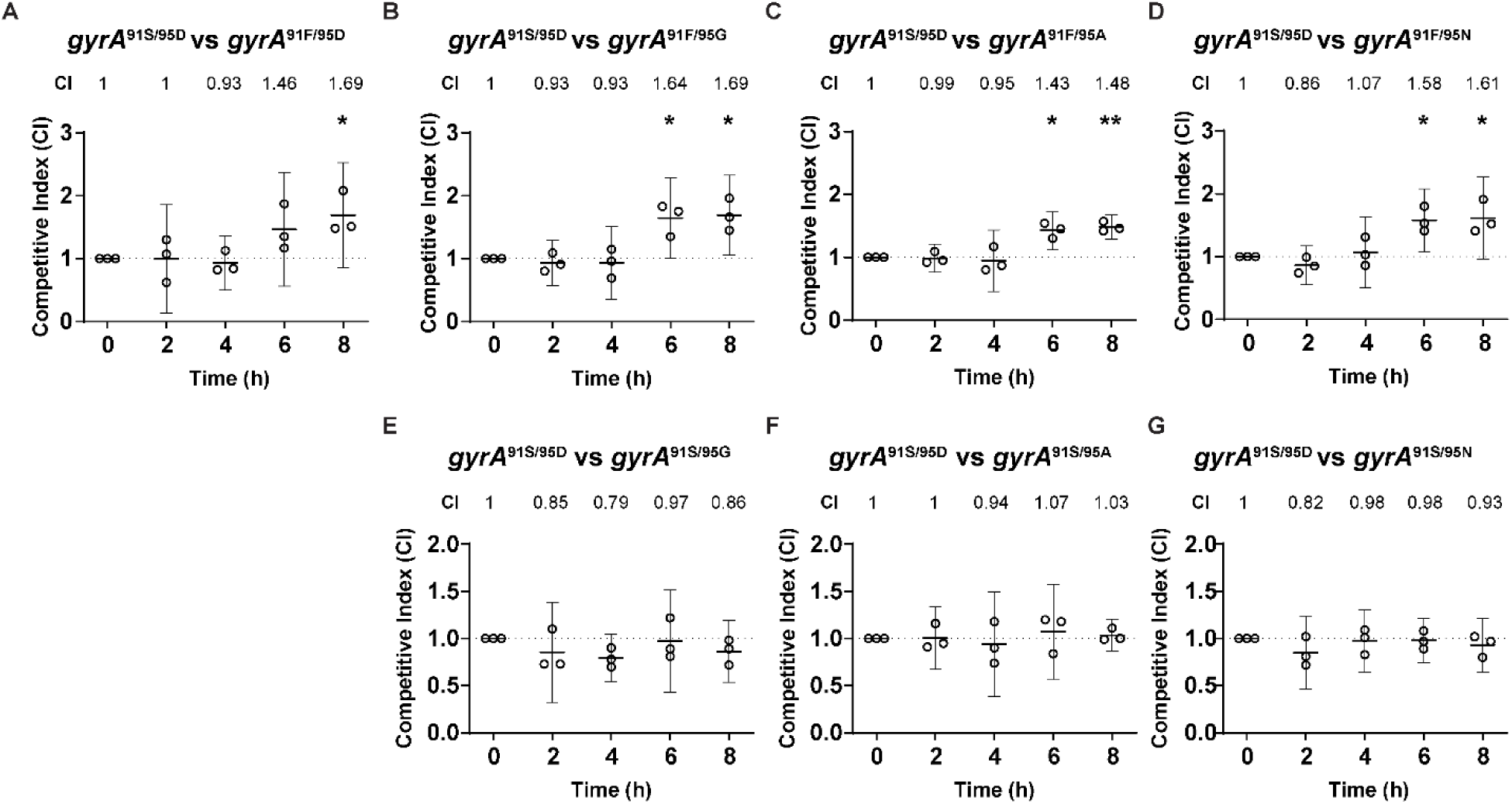
Relative fitness of the *gyrA*^91S/95D^ strain compared to other isogenic *gyrA* mutants. Y-axes show the competitive index (CI) of *gyrA*^91S/95D^ relative to each *gyrA* mutant. The *gyrA*^91S/95D^ strain carried a kanamycin marker and was cocultured with unmarked *gyrA* mutant strains. (A) *gyrA*^91S/95D^ over *gyrA*^91F/95D^: p = 0.96, 0.76, 0.13, 0.04, respectively for 2, 4, 6, and 8 hours. (B) *gyrA*^91S/95D^ over *gyrA*^91F/95G^: p = 0.96, 0.97, 0.02, 0.01, respectively for 2, 4, 6, and 8 hours. (C) *gyrA*^91S/95D^ over *gyrA*^91F/95A^: p = 0.18, 0.27, 0.02, 0.006, respectively for 2, 4, 6, and 8 hours. (D) *gyrA*^91S/95D^ over *gyrA*^91F/95N^: p = 0.2, 0.7, 0.01, 0.02, respectively for 2, 4, 6, and 8 hours. (E) *gyrA*^91S/95D^ over *gyrA*^91S/95G^: p = 0.15, 0.053, 0.35, 0.11, respectively for 2, 4, 6, and 8 hours. (F) *gyrA*^91S/95D^ over *gyrA*^91S/95A^: p = 0.27, 0.69, 0.22, 0.1, respectively for 2, 4, 6, and 8 hours. (G) *gyrA*^91S/95D^ over *gyrA*^91S/95N^: p = 0.23, 0.1, 0.05, 0.22, respectively for 2, 4, 6, and 8 hours. n = 3, representative of three independent experiments performed in absence of any antibiotic pressure. Shown: mean with 95% confidence interval. Statistically significant differences in competitive indices compared to time 0 were analyzed using an unpaired Student’s *t-test*, indicated *p ≤ 0.05 and **p ≤ 0.01.

**Supplementary Figure 7:**
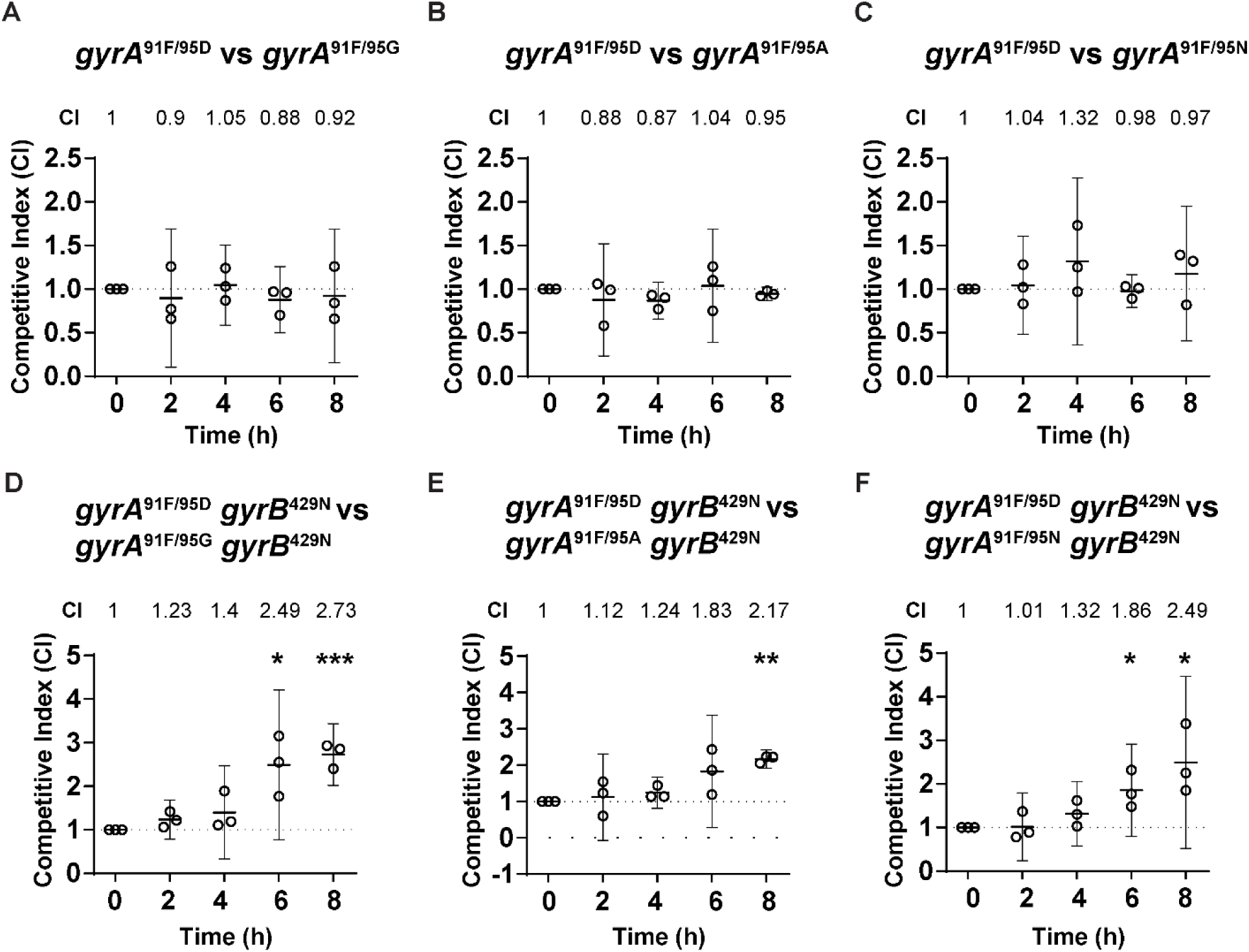
Relative fitness of the *gyrA*^91F/95D^ genotype compared to isogenic *gyrA*^S91F^ strains with various *gyrA*^95^ mutations, alone (A-C) or in combination with the *gyrB*^D429N^ mutation (D-F). (A) *gyrA*^91F/95D^ over *gyrA*^91F/95G^: p = 0.58, 0.84, 0.31, 0.65 respectively for 2, 4, 6, and 8 hours. (B) *gyrA*^91F/95D^ over *gyrA*^91F/95A^: p = 0.35, 0.15, 0.88, 0.31, respectively for 2, 4, 6, and 8 hours. (C) *gyrA*^91F/95D^ over *gyrA*^91F/95N^: p = 0.21, 0.1, 0.1, 0.14, respectively for 2, 4, 6, and 8 hours. (D) *gyrA*^91F/95D^ *gyrB*^429N^ over *gyrA*^91F/95G^ *gyrB*^429N^: p = 0.07, 0.13, 0.02, 0.0006, respectively for 2, 4, 6, and 8 hours. (E) *gyrA*^91F/95D^ *gyrB*^429N^ over *gyrA*^91F/95G^ *gyrB*^429N^: p = 0.62, 0.96, 0.09, 0.002, respectively for 2, 4, 6, and 8 hours. (F) *gyrA*^91F/95D^ *gyrB*^429N^ over *gyrA*^91F/95G^ *gyrB*^429N^: p = 0.7, 0.34, 0.05, 0.04, respectively for 2, 4, 6, and 8 hours. n = 3, representative of three independent experiments performed in absence of any antibiotic pressure. Shown: mean with 95% confidence interval. Statistically significant differences in competitive indices compared to time 0 were analyzed using an unpaired Student’s *t-test*, indicated *p ≤ 0.05, **p ≤ 0.01, and ***p ≤ 0.001.

